# Changes in Neural Dynamics of Brain Activity and Connectivity Independently Predict Post-Stroke Aphasia Recovery

**DOI:** 10.64898/2026.09.17.751646

**Authors:** Andrea Bruera, Zhizhao Jiang, Dorothee Saur, Anika Stockert, Gesa Hartwigsen

## Abstract

Post-stroke language recovery involves reorganization of left-hemisphere language and bilateral domain-general multiple-demand networks. Neuroimaging predictors of recovery are widely used, yet, they are often grouped into broad predictor families, limiting clinical translation and the identification of specific therapeutic targets. Here, we assess brain region-specific neural predictors of recovery using task-related activity and connectivity within a multivariate, cross-validated predictive framework. Forty-seven patients with aphasia were examined longitudinally in the acute, subacute, and chronic phases after ischemic stroke using a sentence comprehension paradigm during functional neuroimaging, with regions of interest spanning language and multiple-demand networks. Confound-controlled multivariate regularized regression with feature ablation tested whether predictor families explained language recovery beyond lesion, age, and aphasia severity, and identified the most relevant region-specific predictors. Early recovery was independently predicted by acute and subacute activity and connectivity, whereas later improvement was predicted by activity alone. Subacute connectivity from multiple-demand to language regions predicted early recovery, while acute multiple-demand network activity predicted long-term outcomes. Later subacute-to-chronic recovery was instead predicted by subacute and chronic activity in bilateral inferior frontal regions. These findings identify phase-specific functional biomarkers of aphasia recovery beyond traditional clinical measures, highlighting multiple-demand and inferior frontal regions as candidate targets for phase-adapted neurostimulation approaches.

## Introduction

Aphasia affects approximately one-third of patients after acute stroke, yet fewer than one-quarter fully recover their language abilities ^1^. Recovery trajectories vary widely ^2^ due to demographic, behavioral, and neural factors, making it difficult to predict outcomes and tailor therapeutic interventions. Although speech and language therapy improves recovery ^3^, maximizing treatment benefit requires accurate prediction of treatment responses ^4,5^, together with a mechanistic understanding of the neural processes that support language recovery over time ^6^.

Aphasia recovery can be characterized in two complementary ways. One is the level of *language ability* achieved at a given phase after stroke, reflecting the patient’s current functional state. The other is the amount of *language improvement* between recovery phases, reflecting recovery-related change over time. Likewise, outcome prediction can serve two distinct purposes: it can explain current language performance or prospectively predict future language ability and improvement from an earlier recovery phase. These complementary prediction trajectories likely reflect partly distinct neural mechanisms, yet they have rarely been examined within a common framework.

Previous studies have identified several robust predictors of aphasia recovery. Initial aphasia severity ^1^ and lesion size ^7^ consistently explain substantial variability in language outcomes, but demographic and other behavioral factors ^1,5^ as well as lesion location have also been associated with aphasia recovery ^7,8^. Beyond these traditional clinical predictors, structural and functional neuroimaging measures – including task-related activity and connectivity – may further increase prognostic accuracy ^9,10,11,12^. Indeed, converging evidence suggests that successful recovery depends on the dynamic reorganization of distributed brain networks, including left-hemisphere language regions, bilateral domain-general multiple-demand regions, and interactions between these systems ^6,13,14,15,16^.

Despite these advances, it remains unclear which neural mechanisms provide the most informative predictions of recovery. Most predictive models integrate multiple *sets* of structural and functional neuroimaging features ^9,12,17^ to maximize prediction accuracy and sensitivity, however, the weight of *individual* predictors and the contribution of specific brain regions and connectivity patterns remains elusive. Moreover, previous work has predominantly focused on resting-state connectivity ^16^, which provides limited cognitive specificity compared to task-related connectivity ^18^, and only few studies have followed patients longitudinally from the acute to the chronic phase after stroke ^10, 16^. As a result, it remains unknown whether activity and task-related connectivity make distinct contributions to predicting current and future (prospective) language recovery across different recovery phases. Identifying *when and where* specific mechanisms predict recovery could help define optimal therapy, targets and time windows, for example for neurostimulation-based interventions.

Here, we addressed these questions using longitudinal task-based functional MRI data from patients with aphasia following first-time acute ischemic stroke. We applied a confound-controlled multivariate predictive framework to investigate how regional task-evoked activity and effective connectivity within and between left-hemispheric language regions, the right inferior frontal cortex, and bilateral multiple-demand network regions contribute to aphasia recovery. Specifically, we distinguished four complementary prediction trajectories: current and prospective prediction of language ability and language improvement. Using cross-validated ridge regression and feature ablation analyses, we evaluated the out-of-sample predictive performance of activity and connectivity for each trajectory. We quantified the independent predictive value of functional neuroimaging beyond traditional clinical predictors and identified the region-specific activity and connectivity patterns that contributed most strongly to prediction. We hypothesized that task-related activity and connectivity would provide complementary predictive information beyond traditional predictors, with multiple-demand regions and right inferior frontal cortex contributing predominantly during early recovery and left-hemispheric language regions becoming increasingly important during later recovery stages.

## Results

We analyzed longitudinal behavioural and task-fMRI data from 47 patients with aphasia, acquired during the acute (≤1 week), subacute (1–3 weeks), and chronic (>3 months) phases after first-ever left-hemispheric ischemic stroke (**SI Table 1** for details). We first evaluated current and prospective prediction of language ability at each time point and language improvement at each time interval (early: acute to subacute, long-term: acute to chronic, late: subacute to chronic) beyond established clinical predictors. We then identified the regional task-evoked activity and effective connectivity measures that contributed most strongly to prediction (**Figure 1**).

**Figure 1.**
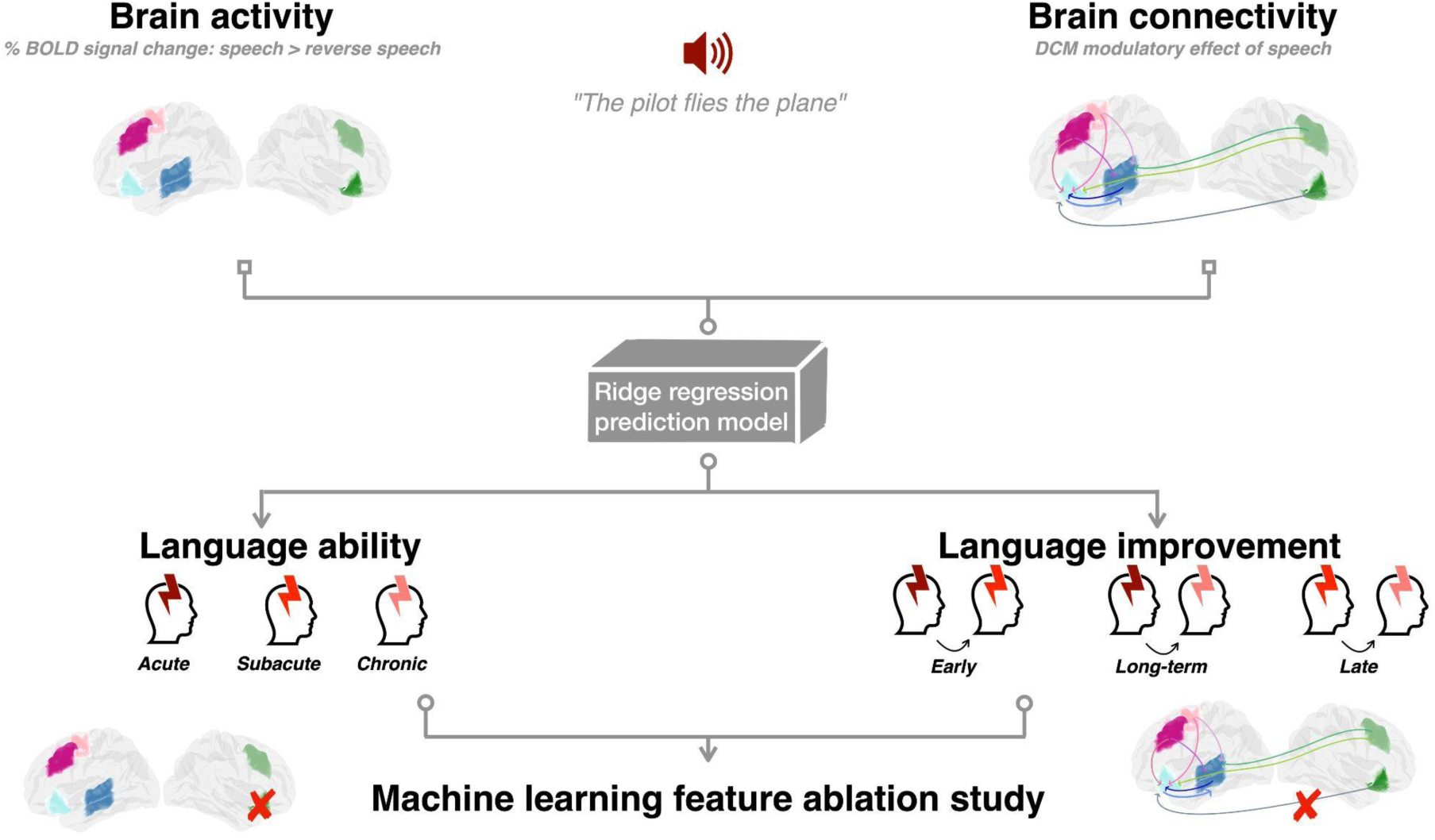
Predictive analysis framework. Longitudinal task-related brain activity (six regions of interest) and effective connectivity (nine connections of interest) were collected from 47 patients with aphasia during the acute, subacute and chronic phases after stroke, using an auditory comprehension task contrasting speech and reversed speech. Cross-validated ridge regression was used to assess to what extent these neuroimaging measures could predict language ability and improvement at each different time point and time interval (early: acute to subacute, long-term: acute to chronic, late: subacute to chronic) as measured with the Aachen Aphasia Test. To assess the unique predictive contribution of functional neuroimaging, prediction targets were residualized for age, lesion affection, previous language ability and the respective other neuroimaging modality before model fitting. Feature ablation was subsequently performed to identify the predictors that contributed most strongly to model performance and to characterize their associations with each prediction target. Abbreviations: BOLD: Blood-Oxygenation-Level Dependent, DCM: Dynamic Causal Modelling.

### Functional neuroimaging predicts recovery beyond traditional predictors

We adopted a cross-validated ridge regression to test whether different functional neuroimaging measures can predict current and prospective language recovery. Comparisons between traditional and functional neuroimaging predictors at each time point and time interval are displayed in **Figure 2**. The reported values refer to the average across all predictors; as statistical tests, we used two-tailed non parametric permutation tests. Note that henceforth, all reported p-values in the results section are corrected for multiple comparisons using the False Discovery Rate (FDR) procedure ^19^.

**Figure 2.**
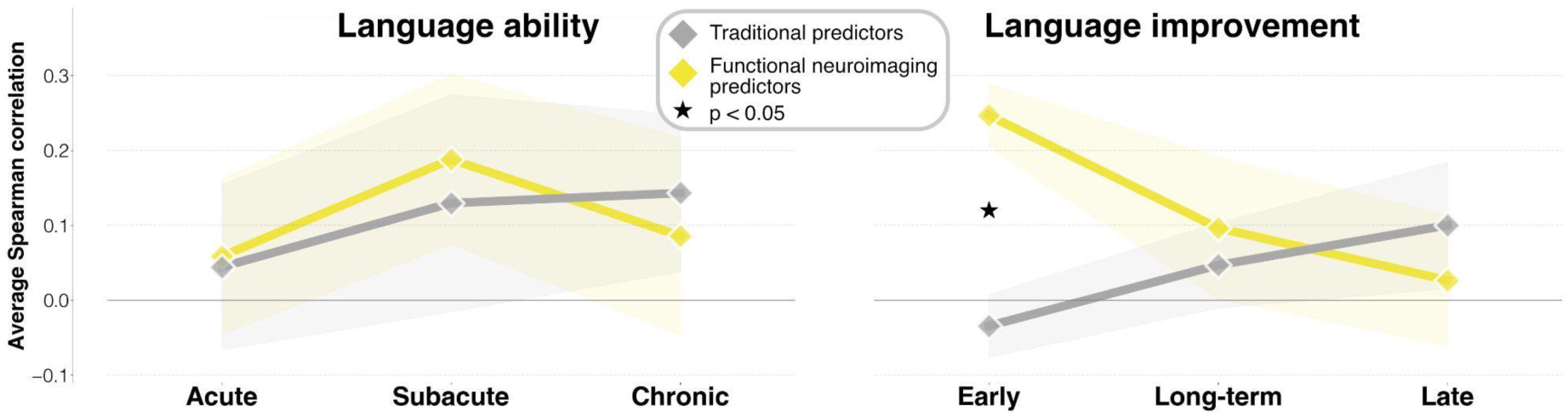
Comparisons between traditional and functional neuroimaging predictors at each time point and interval for language ability and improvement. Each diamond-shaped dot represents the average Spearman correlation (y-axis) between true and predicted targets across 50-iterations of Monte Carlo cross-validated predictions at the corresponding time point (x-axis), for all predictors within each predictor type (traditional predictors: the best predictor among lesion affection, age, previous language ability; functional neuroimaging predictors: task-related brain activity and effective connectivity during speech comprehension). Prediction scores were obtained after removing the variance in targets explained by all other predictors; they therefore refer to the amount of variance explained specifically by the corresponding predictor, above and beyond all other predictors. Lines are added to show the development over time; shaded portions of the plot represent the standard deviation across the individual predictors. Significant differences in predictive performance between traditional and functional neuroimaging predictors are marked with a black star (non-parametric permutation testing, p<0.05, FDR-corrected).

To correctly interpret these results, note that when predicting language ability or improvement from a given predictor (or predictor set), we always first removed the variance explained by all other predictors. Reported prediction scores therefore reflect the unique predictive contribution of the predictor(s) of interest, *over and above the variance already explained by the remaining predictors*. Differences (FDR-corrected) in predictive performance between traditional and functional neuroimaging predictors were significant only for early improvement, where functional predictors provided significantly better predictions (average correlation of functional predictors=0.246, average correlation of traditional predictors=-0.034, statistical comparison (non-parametric permutation test): p=0.0059). At all other time points, functional and traditional predictors had similar predictive performance (functional predictors – acute ability: 0.058; subacute ability: 0.188; chronic ability: 0.085; long-term improvement: 0.096; late improvement: 0.026; traditional predictors – acute ability: 0.044; subacute ability: 0.129; chronic ability: 0.143; long-term improvement: 0.046; late improvement: 0.1; statistical comparisons: acute ability: p=0.949; subacute ability: p=0.74; chronic ability: p=0.728; long-term improvement: p=0.728; late improvement: p=0.671). This suggests that neuroimaging and clinical predictors capture complementary, equally important information.

Zooming in on the individual predictive scores of traditional predictors (**SI Figure 2 and SI Table 2** for details), we found, as expected, that lesion affection could reliably predict language ability at all time points (acute: average correlation=0.263, p=0.0024; subacute: average correlation=0.373, p=0.0024; chronic: average correlation=0.274, p=0.0024), as well as previous language ability (subacute: average correlation=0.214, p=0.0024; chronic: average correlation=0.141, p=0.0085). However, when focusing on language improvement, we found that lesions could only reliably predict long-term (average correlation=0.151, p=0.0024) and late improvement (average correlation=0.205, p=0.0024) – for which also age and previous language ability had above-chance performance (acute ability: average correlation=0.141, p=0.0057; age: average correlation=0.154, p=0.0042).

Next, we focused on the predictive performance of models trained on each family of predictors (functional activity or connectivity) at different time points (**Figure 3**). Prediction scores represent the unique variance explained by each family of predictors after accounting for established traditional predictors (age, lesion affection, and aphasia severity) and the complementary functional neuroimaging measure acquired at the same time point (e.g., when evaluating acute effective connectivity, variance explained by acute task-evoked activity was removed). Prediction performance for traditional predictors and complementary functional neuroimaging predictors (confound control) is reported in **SI Figure 2** and **SI Table 2**. Most models predicted language outcomes significantly above chance.

**Figure 3.**
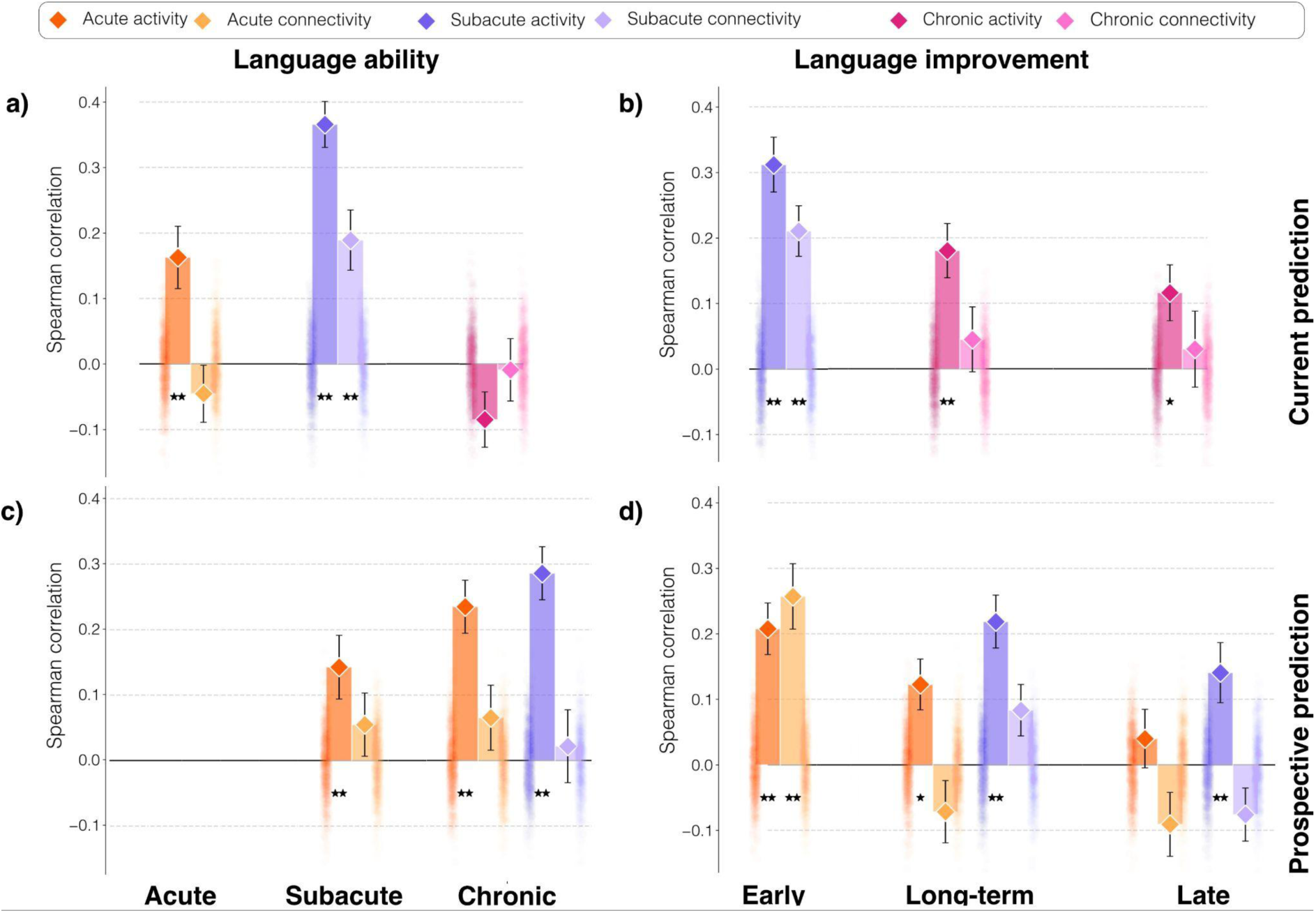
Ridge regression prediction scores for current and prospective prediction of language ability and improvement. Bars represents the average Spearman correlation between true and predicted targets across 50-iterations of a Monte Carlo cross-validated prediction. Prediction scores were obtained after removing the variance in targets explained by the other family of neuroimaging predictors, lesion affection, age and – for language improvement (right panels) – previous aphasia severity. Results for the null distribution obtained with 1000 random permutations are displayed as individual scatter points next to the corresponding bar. Highlighted similarities are significantly above chance (*/**p < 0.05/0.005, FDR-corrected). Left panels (a, c) show the ability of neuroimaging predictor sets (brain activity and connectivity) to predict language ability during each phase. Right panels (b, d) show the extent to which the same sets of variables can predict language improvement from the acute to subacute (early), the acute to chronic (long-term) and the subacute to the chronic phases (late), respectively. For sets significantly predicting targets (*/**), post-hoc ablation analyses were performed.

### Functional neuroimaging predicts current recovery

When predicting ***language ability*** at the same time point of the functional neuroimaging measurements (current prediction; **Figure 3a**), we found that acute ability could be predicted only by acute activity (average correlation=0.163, p=0.0028), while subacute ability could be predicted by both subacute activity (average correlation=0.366, p=0.0028) and connectivity (average correlation=0.189, p=0.0043).

For predictions of ***language improvement*,** we found a greater predictive role of connectivity measures, specifically for early improvement (from acute to subacute). For predictions at the same time point of the measurements (current predictions; **Figure 3b**), both subacute activity and connectivity predicted significantly better than chance early improvement (activity: average correlation=0.312, p=0.0028; connectivity: average correlation=0.211, p=0.0028); long-term and late improvement, by contrast, could be predicted only by chronic activity (long-term: average correlation=0.18, p=0.0028; late: average correlation=0.116, p=0.016) and not by chronic connectivity (long-term: average correlation=0.045, p=0.2267; late: average correlation=0.03, p=0.3823).

### Functional neuroimaging predicts future recovery

When predicting ***future language ability*** (prospective prediction; **Figure 3c**) we found that only activity could reliably do so: subacute ability could be predicted by acute activity (average correlation=0.143, p=0.0043) and chronic ability by both acute (average correlation=0.235, p=0.0028) and subacute activity (average correlation=0.286, p=0.0028).

For ***prospective language improvement*** (prospective prediction; **Figure 3d**), early improvement could be predicted by both acute activity and connectivity (activity: average correlation=0.208, p=0.0028; connectivity: average correlation=0.257, p=0.0028). Acute and subacute activity predicted long-term improvement significantly better than chance (acute: average correlation=0.123, p=0.0168; subacute: average correlation=0.219, p=0.0028), and subacute activity was the only reliable neuroimaging predictor of late improvement (average correlation=0.141, p=0.0043).

### Identifying the most important individual neuroimaging predictors of language ability and improvement

Beyond identifying sets of neuroimaging predictors that reliably predicted language ability and improvement in the main analyses, we aimed to determine which individual functional neuroimaging measures contributed most to predicting language recovery and their relationship with behavior. Using feature ablation analyses, we explored the contribution of ROI-specific predictors of functional activity and connectivity, only for prediction models where prediction was significantly above chance in the main analyses (p<0.05 FDR corrected). We report the individual predictors whose exclusion had a significant impact on the regression model’s ability to learn the target variables, analyzed separately for each predictor set at the corresponding time points (**SI Tables 4-5** for complete statistics). A significant impact was accepted if the model’s prediction performance was no longer significantly above chance, as indicated by the p-values after removal reported below. The average model weight of a given predictor indicated the direction of the relationship, with positive or negative weights reflecting that higher activity or connectivity values were associated with more or less language ability or improvement, respectively.

### Individual brain activity predictors

Acute activity in the left DLPFC was the individual ROI with the strongest, significant impact on predictions of *subacute* (p after removal=0.2927) and *chronic language ability* (p=0.0957) (**Figure 4**, left). A positive relationship indicated that higher activity predicted better language ability in the respective phase (subacute: average weight=0.0531; chronic: average weight=0.0382). Acute activity in the left SMA also caused a significant decrease in prediction performance (p=0.0522) of *acute language ability*, again higher activity predicted better ability (average weight=0.0308).

**Figure 4.**
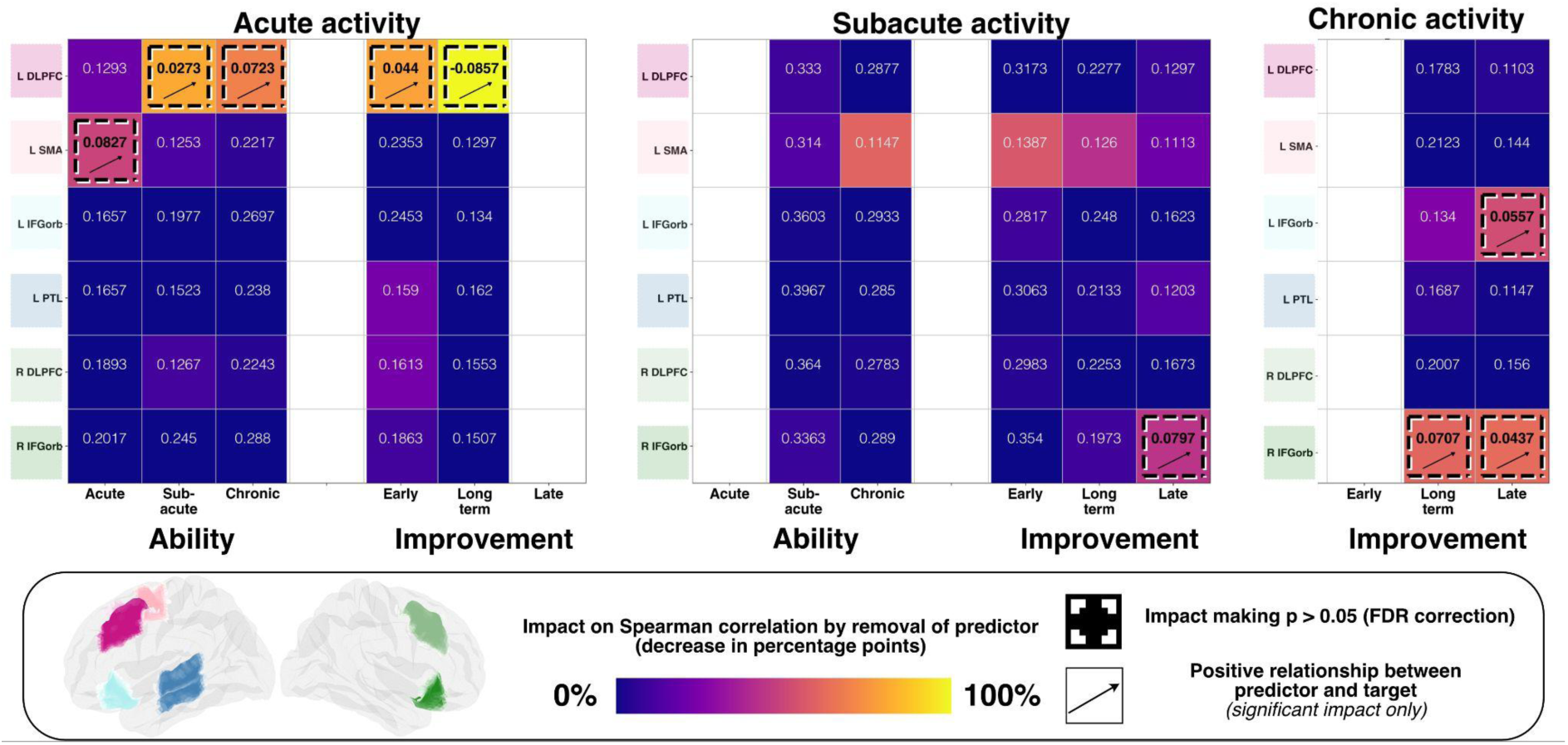
Impact and directionality of phase-specific activity in individual ROIs on model predictions of language ability and improvement. The plot is divided into three subsections, one for each predictor and time point. Each heatmap cell represents the predictive performance (Spearman correlation), after removing the corresponding ROI activity of the respective phase from the predictor set. Cell colours reflect the percentage of decrease in predictive performance. Dashed squares indicate predictive performance of the remaining predictors to be at chance (i.e. not significant, with p > 0.05, FDR corrected) after the individual predictor was removed. Abbreviations: L: left, R: right, DLPFC: dorsolateral prefrontal cortex, SMA: supplementary motor area, IFGorb: pars orbitalis of the inferior frontal gyrus, PTL: posterior temporal lobe.

Acute activity in the left DLPFC also had a significant impact on prediction of *early language improvement* (p=0.2081) with higher activity predicting greater language gains (average weight=0.0522). Acute activity in the left DLPFC (p=0.955) and chronic activity in the right IFG (p=0.088) were the most important individual predictors for long-term improvement, again showing a positive relationship (acute DLPFC: average weight=0.0144; chronic right IFG: average weight=0.018). For *late improvement*, subacute and chronic activity in the right IFG were the most relevant predictors (p=0.0604 and 0.1853) (**Figure 4**, middle and right) and were positively related to behavioral gain (average weight=0.0144 and 0.003). The same was true for chronic activity in the left IFG (p=0.1202, average weight=0.0051).

### Individual brain connectivity predictors

Connectivity from left DLPFC to left IFG in the subacute phase was the most important predictor of *subacute language ability* (p=0.2773) (**Figure 5**, left) and of *early language improvement* (p=0.1841) (**Figure 5**, right), in both cases, higher facilitatory effective connectivity was positively related to subacute ability (average weight=0.1276) and early improvement (average weight=0.1144). No other connectivity measure had, individually, a significant effect on prediction scores. Complete statistics can be found in the Supplementary Information.

**Figure 5.**
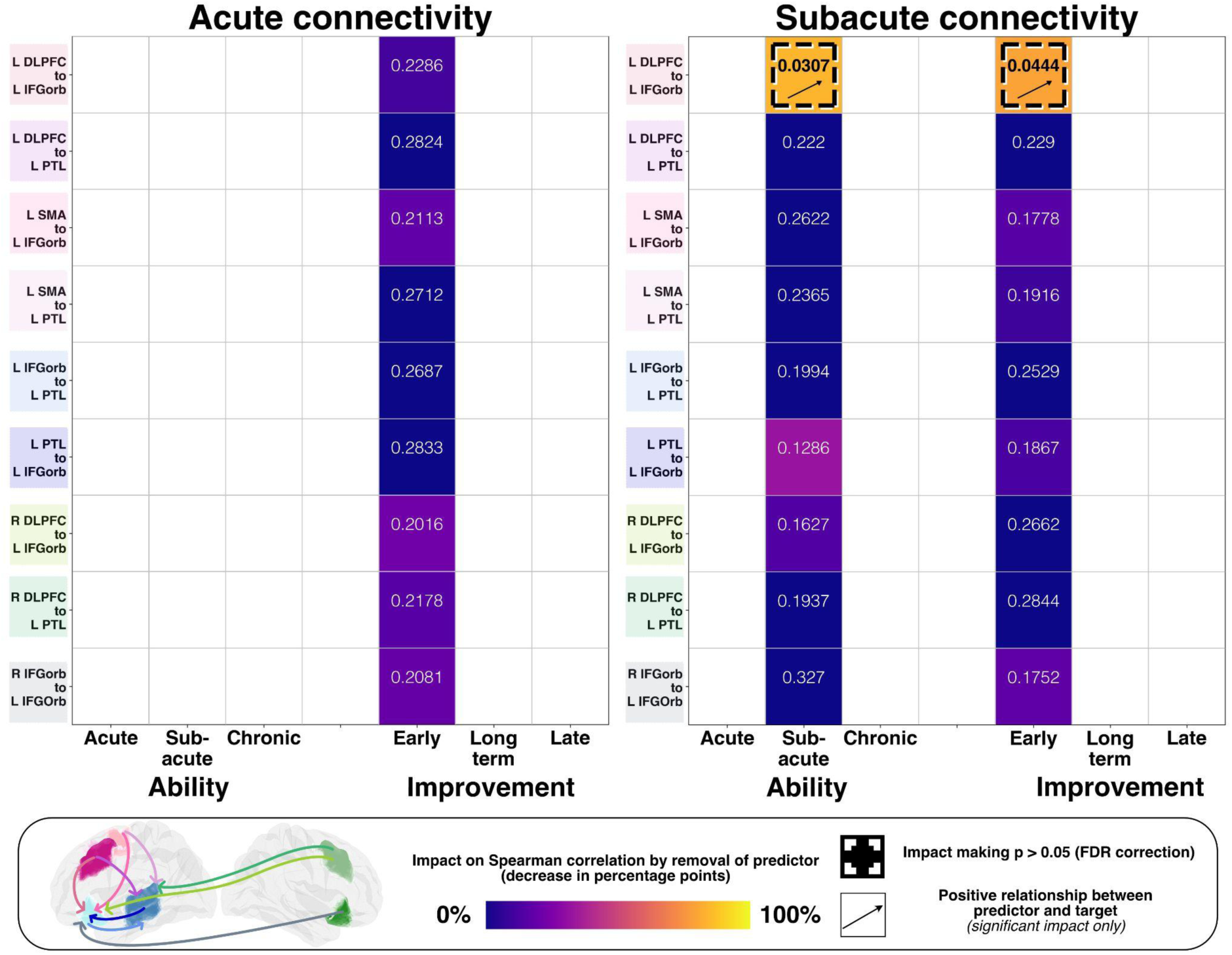
Impact and directionality of acute and subacute connectivity in individual ROIs on model predictions of early, long-term and late language ability and improvement. The plot is divided in two subsections, one for each predictor and time point. Each heatmap cell represents the predictive performance (Spearman correlation) after removing the corresponding connectivity measure of the respective phase from the predictor set. Dashed squares indicate predictive performance of the remaining predictors to be at chance (i.e. not significant, with p > 0.05, FDR corrected) after the individual predictor was removed. Abbreviations: L: left, R: right, DLPFC: dorsolateral prefrontal cortex, SMA: supplementary motor area, IFGorb: pars orbitalis of the inferior frontal gyrus, PTL: posterior temporal lobe.

## Discussion

We used machine learning to evaluate longitudinal functional neuroimaging as a predictor of aphasia recovery. We assessed which predictor families – task-related activity or connectivity, at different recovery phases – predicted language ability and improvement after adjusting for traditional predictors (age, lesion affection and aphasia severity) and the respective other predictor family. Using feature ablation, we identified the most important individual predictors. To our knowledge, this is the first study to systematically demonstrate the independent and phase-specific contribution of longitudinal task-related activity and connectivity as functional biomarkers of aphasia recovery beyond established traditional predictors. Spanning the acute, subacute and chronic phases after ischemic stroke, our findings identify *whether, when* and *which* patterns of functional brain activity and effective connectivity provide information of current function and future recovery potential.

Three main findings emerged. First, functional neuroimaging predicted aphasia recovery independently of established traditional predictors, with the strongest contribution during early recovery, from the acute to subacute phase, when trajectories are highly dynamic and prognosis is challenging ^1^ . Specifically, activity and connectivity enabled both current and prospective prediction of early improvement, whereas traditional predictors alone showed no significant predictive value. In contrast, for language ability and later improvement, functional neuroimaging and traditional predictors contributed comparably but independently (**Figure 2**), suggesting that they capture complementary rather than redundant aspects of recovery. This pattern helps reconcile discrepant findings in the literature: Whereas several studies concluded that lesion characteristics and baseline aphasia severity account for most variance in recovery ^20,21,22^, with functional imaging (resting-state connectivity or task-related activity) adding little incremental value ^23^, others demonstrated improved prediction when chronic task-related activity or normative sample functional connectivity were incorporated ^24^. These discrepancies likely reflect differences in recovery phase, imaging modality, prediction target, and whether imaging was evaluated independently or alongside traditional predictors. More fundamentally, our findings suggest that traditional predictors and functional biomarkers capture complementary biological processes: lesion characteristics describe the structural consequence of stroke, whereas functional biomarkers capture the dynamic physiological state of the remaining undamaged brain ^25^. By systematically comparing these predictor classes across recovery phases within the same longitudinal cohort, we further show that their relative contribution shifts over time, with functional biomarkers providing the strongest unique contribution early after stroke. This likely reflects transient processes – edema, altered cerebral perfusion, and diaschisis-related disconnection in structurally intact regions ^26, 27^ – that shape early language recovery but are not captured by lesion characteristics alone. As recovery progresses, the contribution of functional biomarkers and traditional predictors becomes more balanced: while structural damage and age likely constrain the ceiling of later recovery ^28,29^, functional biomarkers continue to reflect the ongoing reorganization of residual language networks. Collectively, these findings indicate that aphasia recovery depends not only on the integrity of preserved brain regions but also on the capacity of distributed brain networks to flexibly reorganize over time ^16, 30, 31^.

Our second finding shows that the identified functional biomarkers predict both current and forecast future recovery (**Figure 3**). Measures acquired at a given recovery phase predicted language ability within that same phase or at the end of an interval, indicating that task-related activity – and during the subacute phase, effective connectivity – closely reflect the current functional capacity of the underlying brain networks. These measures can therefore be conceptualized as *functional state biomarkers*. Likewise, measures acquired during an earlier phase predicted language ability or improvement at a later phase, capturing information about the brain’s future recovery potential; we conceptualize these as *functional prognostic biomarkers*. Importantly, prognostic biomarkers reflect the biological capacity for recovery rather than the recovery ultimately achieved, which is also shaped by other environmental and therapeutic factors, such as social participation and the intensity of speech and language therapy ^3^, that were not explicitly modelled in the present study. Incorporating such variables alongside functional neuroimaging may further improve prediction accuracy.

Prospective prediction varied by recovery phase. Task-related activity from the acute and subacute phases predicted subsequent language ability, indicating that preserved or compensatory recruitment of brain regions early after stroke reflects the capacity of undamaged brain networks to support later outcome. Early improvement (acute to subacute) was independently predicted by both acute activity and effective connectivity, whereas later improvement (subacute to chronic) was predicted primarily by subacute activity. Notably, subacute measures provided the most substantial predictions of both language improvement and later ability, likely reflecting a stabilization of post-stroke physiology alongside the emergence of meaningful network reorganization. In contrast, acute measures may be influenced by transient disturbances in cerebral physiology ^32^, while chronic-phase patterns may become more heterogeneous because patients achieve recovery through distinct compensatory network configurations partly constrained by lesion location ^14, 15^. Collectively, our finding suggests that activity and effective connectivity capture complementary aspects of recovery that shift across stages, with the subacute phase representing a critical window in which regional activity and network interactions jointly shape the trajectory of subsequent recovery.

As a third finding, we quantified the contribution of specific activity and connectivity patterns to predicting recovery over time. Feature-ablation analyses revealed that the relative contribution of the multiple-demand and language networks to recovery changes systematically over time. Acute (and subacute) task-related activity within multiple-demand network regions (left DLPFC, SMA) emerged as the most important predictor of language ability across all recovery stages and of both early and long-term improvement. Since the multiple-demand network supports domain-general cognitive control and adaptive task performance ^33^, preserved or compensatory early recruitment of these regions may provide a general neural resource enabling residual function despite widespread disruption of the language network. Rather than reflecting language-specific processing, activity within this network likely indexes the capacity of the injured brain to engage residual neural resources during the acute and subacute phases. In contrast, later language improvement was most strongly predicted by left and right IFG activity during the subacute and chronic phases, indicating that the neural mechanisms supporting recovery shift over time. This is congruent with previous work emphasizing an early role of the multiple-demand network ^34^ and a later normalization of activity in language network regions ^13, 14^. It also supports both chronic right hemisphere involvement in some patients ^35^ and a later activity shift towards the language network being beneficial for language recovery and successful therapy ^36,37^.

This result substantially extends previous studies by revealing a complementary role of activity and connectivity. No single acute connection predicted early (acute to subacute) improvement, suggesting the extent of widespread dysfunction (i.e., diaschisis) rather than affection of a specific system contributing to early recovery ^26^. In contrast, early improvement and subacute ability were predicted not only by increased activity, but also by subacute connectivity from multiple-demand to language regions (left DLPFC to left IFG), as most relevant functional state and prognostic biomarker. Together, this suggests that recovery not only depends on (re-)activation, but also on the interaction between multiple-demand and language network regions, particularly in the subacute phase, followed by increased reliance on activity in language regions and the homologous right IFG, as recovery progresses. This extends the previously suggested early contribution of the right IFG during the subacute phase after stroke to a more sustained role later in recovery ^13^.

The main findings for predictors of language improvement are summarized in **Figure 6**.

**Figure 6.**
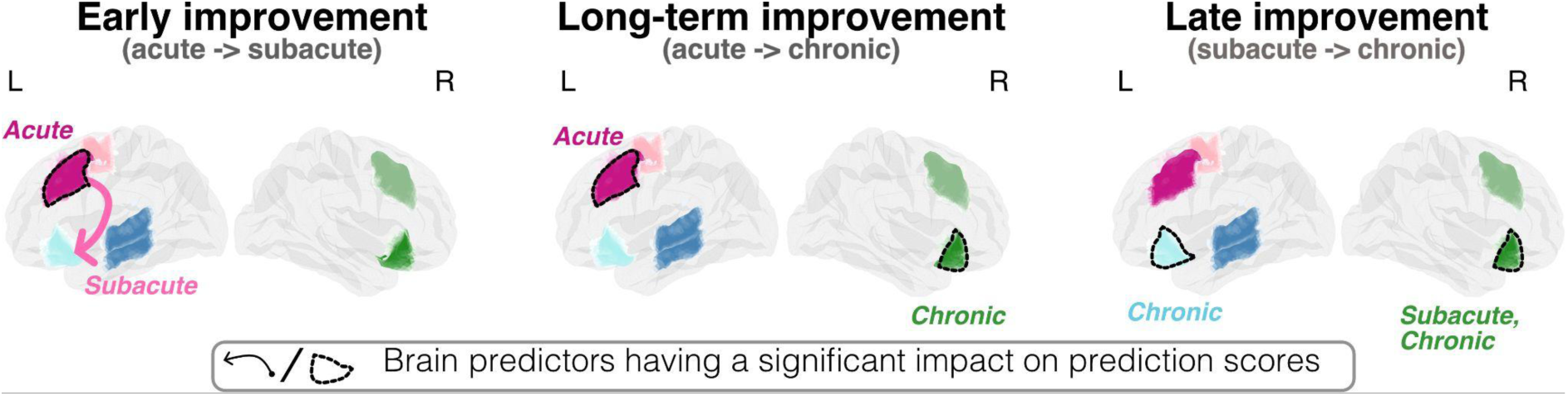
Summary of the main findings for predictors of language improvement after stroke. Schematic visualization of the most important predictor regions or connections from each predictor family (brain activity and effective connectivity) that was found to be a significant predictor of language improvement in the main analyses. Significant ROIs for brain activity are marked by thick black lines. Connectivity measures are indicated by an arrow, whose tip indicates the direction. Phases for the corresponding measure are visualized next to each measure.

Identifying not only *whether and when* functional neuroimaging predicts aphasia recovery, but also *which* neural systems are most informative may help inform the selection and timing of therapeutic interventions. One important implication is that neurostimulation approaches could benefit from dynamic, phase-specific protocols targeting different nodes across the course of aphasia recovery ^38^, rather than a single cortical target, to support the evolving network mechanisms underlying recovery. This network-oriented perspective is consistent with previous findings in healthy volunteers that stimulation of one cortical region can modulate effective connectivity in distributed language networks and produces behaviorally relevant after-effects on language comprehension ^39^. Testing whether phase-specific stimulation targets improve recovery outcomes will require dedicated interventional studies.

Most previous neurostimulation studies in aphasia have targeted the language network or its right hemisphere homologues in the chronic phases of recovery using varying facilitatory and inhibitory protocols ^40^. Our results motivate extending such approaches to earlier phases (i.e., the first weeks) after stroke and targeting regions and functional interactions that are critical during the respective recovery period, as they likely represent a critical window and substrate of heightened reorganization where therapeutic interventions could be particularly effective. Given that early recovery was strongly related to the recruitment of multiple demand network regions and their integration with residual language-network regions, early interventions – as an adjunct to language and cognitive training – may benefit from facilitating activity of multiple demand network regions, such as the left DLPFC, while strengthening integration with undamaged language network regions such as the left IFG. This is consistent with initial evidence that stimulation of personalized target regions outside the language network during aphasia therapy can improve language functions and modulates functional connectivity ^41,42^, supporting a network-oriented approach in which stimulation of multiple-demand network regions may enhance recovery through their interactions with the language network.

Our results further suggest that later interventions may progressively shift towards enhancing activity within undamaged language regions, particularly the left and right anterior IFG, at least for supporting recovery of language comprehension as assessed here. Our finding that right anterior IFG activity positively predicted recovery, together with previous evidence of increased IFG activity ^13,14,43^ and enhanced right-to-left IFG connectivity ^15^, suggests a beneficial contribution of the right IFG within the residual network. This contrasts with studies advocating inhibitory right IFG stimulation during later stages of recovery to reduce maladaptive plasticity and shift activity back to preserved left-hemispheric language regions (e.g., ^44^). Such discrepancies likely reflect functional heterogeneity within the right IFG ^45^, limiting our interpretation to a supportive role of its anterior subregion (pars orbitalis) in language comprehension. Future work should directly test this hypothesis and dissociate contributions of different IFG subregions to aphasia recovery.

Our analyses identified phase-specific contributions of multiple demand and inferior frontal regions, but these findings should be interpreted within the constraints of our hypothesis-driven selection of regions and connections, chosen a priori to maximize interpretability and reduce model complexity. Other regions supporting language processing may also be informative but were not evaluated here and should be integrated in future whole-brain or data-driven approaches to identify additional functional biomarkers. In addition, before dynamic approaches or decision support for individualized treatment can be translated into clinical practice, external validation is essential. Although predictive performance was evaluated using out-of-sample cross-validation, models were developed in a single cohort comprising patients with heterogeneous lesion locations and impairment. Different populations or lesion distributions may alter the relative importance of individual biomarkers. Consequently, the present findings should be viewed as initial evidence for the biological and predictive value of longitudinal functional neuroimaging rather than an independently validated clinical prediction model.

In conclusion, this study complements and extends over two decades of longitudinal functional neuroimaging studies on language recovery (e.g., ^13,14,15,46,47^). Beyond identifying which predictors are most informative across the course of aphasia recovery, our findings point toward a route for clinical translation. While lesion characteristics and demographics constrain the limits of later recovery imposed by structural damage or age, functional neuroimaging biomarkers capture the complementary dynamics of cortical reorganization underlying longitudinal aphasia recovery, motivating dynamic, phase- and network-specific neurostimulation strategies tailored to individual patterns of network dysfunction. Although candidate targets and individualized applications require independent prospective validation, our findings illustrate how longitudinal functional neuroimaging could inform precision rehabilitation by guiding phase-specific therapeutic strategies and individualized prediction of post-stroke aphasia recovery.

## Methods

### Participants

We reanalyzed a subset of the dataset reported in ^26^. The sample comprised 47 patients with longitudinal behavioural and neuroimaging data acquired during the acute (≤ 1 week), subacute (1-3 weeks) and chronic (>3 months) phases after first-ever left-hemispheric ischemic stroke resulting in aphasia. To reduce heterogeneity in recovery mechanisms, the chronic assessment window was restricted to the early chronic phase (3-9 months). This restriction follows evidence that most language recovery occurs until the first three months, with comparatively limited gains thereafter ^16^. Around this period, patients are assumed to have completed inpatient and first rehabilitation treatment driven recovery, whereas later improvements are more likely to reflect additional compensatory or outpatient treatment-related mechanisms. Characterizing these later processes is therefore of particular interest, as they may contribute targets for subsequent rehabilitation interventions.

The final sample (mean age 55.2 ± 14.0 years; 13 female) is described in detail in Supplementary Information (SI) **Table 1** and **SI Figure 1**.

Language was assessed at each time point using the Aachen Aphasia Test (AAT ^48^). A composite language comprehension score was derived from the Token Test, the auditory and written comprehension AAT subtests. Scores were scaled between 0 and 1 (1 = unimpaired). This measure reflects acute, subacute and chronic language comprehension performance and is taken as a proxy of aphasia severity in the acute and subacute phases. Hereafter, this variable is referred to as ‘language ability’. Mean language ability increased from 0.48 (interquartile range, IQR 0.52) in the acute phase to 0.65 (IQR 0.49) in the subacute phase and 0.84 (IQR 0.21) in the chronic phase, indicating recovery over time.

### fMRI Design and Analyses

Two similar versions of an event-related functional MRI sentence comprehension paradigm were used (cf. ^10,13^). Participants listened to short German sentences (intelligible speech: e.g. “The pilot flies the plane”) and temporally reversed versions of the same stimuli (unintelligible reversed speech). To ensure attentive listening, they were instructed to press a button. For more details, please refer to the Supplementary Information. A detailed description of MRI acquisition and preprocessing is provided in the methods section of ^15^.

### Region-of-interest (ROI) Selection

ROIs were selected following our previous fMRI studies on longitudinal activity or effective connectivity changes after stroke ^14,15^. The six representative ROIs are also well-supported in the literature of language and cognitive brain networks ^49,50,51^ . For each ROI, a 20 mm spherical mask was created around the Montreal Neurological Institute (MNI) coordinates reported in ^15^. To account for inter-individual variability of peak activation, subject-specific ROIs were defined as the top 10% most activated voxels within each sphere (cf. ^52^).

The left pars orbitalis in the inferior frontal gyrus (IFG) and the posterior temporal lobe (PTL) were assigned to the language network; bilateral dorsolateral prefrontal cortex (DLPFC), and the supplementary motor area / dorsal anterior cingulate cortex (henceforth: SMA) were assigned to the multiple-demand network; and the right IFG represented the right-hemispheric lesion homologue of the left inferior frontal gyrus. Based on our previous studies ^14,15^, nine connections of interest were defined; two connections within the language network: left PTL to IFG, and left IFG to PTL; one connection from right to left IFG, and six connections between the multiple-demand and the language network: right DLPFC to left IFG, right DLPFC to left PTL, left DLPFC to left IFG, left DLPFC to left PTL, left SMA to left IFG, and left SMA to left PTL.

Lesioned and perilesional voxels (0-3 mm beyond the lesion borders) were excluded individually based on the normalized lesions (cf. ^15^) to avoid potential effects of abnormal BOLD signal. Lesion affection per left hemisphere ROI was calculated as the percentage of voxels damaged within each of the four 20 mm spheres. Further details on task-related fMRI and effective connectivity analyses are provided in the Supplementary Information.

### Confound-controlled Predictive Modeling and Feature Ablation

We used a cross-validated predictive framework to investigate how multimodal neuroimaging measures predict the course of post-stroke aphasia recovery. Two complementary prediction settings were considered: (a) *current prediction* of observed language ability and improvement from neuroimaging of the same recovery phase or at the end of an interval and (b) *prospective prediction* of future ability and improvement from earlier neuroimaging measures. Together, these analyses distinguish neural correlates related to achieved recovery and prospective functional biomarkers of future recovery. Ridge regression with L2 regularization and permutation testing was applied to ensure stability and robustness of estimates ^53^. Ridge regression can deal with multicollinearity in the predictors ^54^ – an important factor to consider for our functional neuroimaging-based predictors, because of the physical properties of the BOLD signal across the brain ^55^. L2 regularization was chosen to control for predictor collinearity, as the analysis focused on a limited number of predefined predictors (6 ROIs for activity, 9 for connectivity) rather than high-dimensional voxelwise data.

We used a Monte Carlo cross-validation approach ^56^ with 50 train/test splits (80% training, 20% test). A ridge regression model was trained on the training set (data from 80% of the subjects) and fitted with cross-validated alpha parameter selection (possible alpha between 10^-5^ and 10^4^). The test set (the left-out data from 20% of the subjects) was used to evaluate predictive performance of the model based on the average correlation scores between predicted and observed targets across the Monte Carlo iterations, verifying whether predictions were significantly and reliably better than chance (cf. ^9^).

We repeated the training/testing cross-validation procedure for each set of functional and traditional neuroimaging predictors. Before training each ridge regression model, we first removed the variance explained by all other predictors using the cross-validated confound removal procedure of ^28^, which essentially trains a linear confound removal model on the training set and then removes the variance explained by the confounds from the test set (so-called residualization), before the predictions of interest are computed (for details, see Supplementary Information). This allowed us to assess the unique predictive performance of each set of predictors, beyond what can be explained by all others. To identify which individual predictors contributed most strongly to the predictions, we performed post-hoc analyses for sets of predictors (e.g. brain activity during the acute phase) whose ability to predict a target (e.g. early language improvement) was significantly better than chance (p < 0.05). To obtain measurements of the individual contribution of each predictor, we ran a feature ablation study, a common choice when interpreting machine learning predictive models ^57^: we repeated the analyses after removing each individual predictor and observed the impact of the removal on the average prediction score. An individual predictor was considered to significantly contribute to prediction, if prediction after removal was no longer significant. Here, we used the same cross-validated confound removal procedure to remove the variance explained by other variables.

To interpret the directionality of the predictors (i.e. whether the relation between a specific brain activity or connectivity and the target was direct or inverse), we looked at the average of the ridge regression weight for each predictor across the cross-validation folds. A positive average weight would indicate a direct relationship (an increase in the predictor value leads to an increase in the target value, and *vice versa*), whereas a negative average weight would indicate the opposite (an increase in the predictor value leads to decrease in the target value).

#### Predictors and Targets

Predictors comprised mean language-related activity in six ROIs and effective connectivity in nine connections of interest, measured during acute, subacute and chronic phases. Unlike previous predictive studies using resting-state fMRI, we employed task-based *functional* measures, to capture language-related neural processes.

Out-of-sample prediction analyses were grouped into two settings according to the temporal relationship between predictors and targets. Current prediction used predictors and language ability or improvement obtained during the same phase or at the end of an interval. Prospective predictions used predictors measured at a previous phase or at the beginning of an interval to forecast future ability or improvement (i.e. subacute activity predicting late improvement or chronic ability). Reverse temporal predictions (e.g. subacute activity predicting acute ability) were not performed because they lack biological and clinical interpretability.

Two target variables were modeled: language ability (three measurements: acute, subacute and chronic) and language improvement (three intervals - early: acute to subacute; long-term: acute to chronic; late: subacute to chronic). Language improvement was defined as residualized change after accounting for ability at the beginning of the respective phase, using the cross-validated confound removal described below. Thus, each participant contributed three language ability and three improvement scores. By including predictors and targets with both different spatial and temporal information, the analyses could not only characterize *where*, but also about *when* a certain neuroimaging measure became predictive of recovery.

#### Additional predictors

Age and lesion affection have been found to reliably explain language recovery in post-stroke aphasia ^9,11^. To ensure that our results would not reflect spurious correlation with these variables, we removed the variance that could be explained by lesion affection and age from the targets. In the Results section, we refer to lesion affection, age and previous language ability as *traditional predictors* – as opposed to *functional neuroimaging* predictors (brain activity and effective connectivity), that constitute the novelty of our approach.

We also re-ran the analyses for the traditional predictors, since this allowed us to better understand the relative predictive value of neuroimaging predictors. Results are reported and discussed both in the main text (**Figure 2**; average results) and in the Supplementary Information, where the full set of results for the traditional predictors is reported in **SI Figure 2** and **SI Table 2**.

#### Statistical Testing

The main analyses (**Figure 3**) tested whether each predictor set achieved above-chance performance for the corresponding prediction setting (current and prospective prediction), quantified as the Spearman correlation between predicted and observed target values. Statistical significance was assessed using non-parametric permutation testing for predictive models ^53^. Specifically, the target variables were randomly permuted 1000 times, and the train/test procedure was repeated for each permutation to obtain an empirical null distribution of correlation values under the null hypothesis of no association. One-sided p-values were computed as the proportion of permutations in which the permuted correlation was higher than the observed correlation. To correct for multiple comparisons across predictors sets and time points, p-values were adjusted with the False Discovery Rate (FDR ^19^) procedure and thresholded at p<0.05.

In the feature ablation study (**Figures 4-5**), the same permutation-based framework was applied. Prediction performance was recomputed after removing each predictor in turn and compared against the same null distribution derived from the main analyses. An individual predictor was considered critical for prediction if its removal reduced model performance to a level that was no longer significantly above chance (p < 0.05, FDR-corrected).

## Acknowledgments

GH was supported by the Lise Meitner Excellence Program of the Max Planck Society, the European Research Council (ERC-2021-COG 101043747) and the German Research Foundation (DFG, HA 6314/4-2, Research Unit 5429/1 (467143400), HA 6314/10-1). AB would like to thank Ajay Halai for extensive and insightful feedback regarding an earlier version of these results.

**Author Contributions** (**CRediT system:** Conceptualization, Data curation, Formal analysis, Funding acquisition, Investigation, Methodology, Project administration, Resources, Software, Supervision, Validation, Visualization, Writing – original draft, Writing – review & editing) **AB:** Conceptualization, Data curation, Formal analysis, Investigation, Methodology, Resources, Software, Validation, Visualization, Writing – original draft, Writing – review & editing; **ZJ:** Conceptualization, Methodology, Data curation, Formal analysis, Resources; **DS:** Supervision, Writing – review & editing; **AS:** Conceptualization, Data curation, Methodology, Project administration, Supervision, Writing – original draft, Writing – review & editing; **GH:** Conceptualization, Funding acquisition, Methodology, Project administration, Resources, Supervision, Writing – original draft, Writing – review & editing.

## Data availability

The dataset cannot be shared due to privacy reasons. The code, as well as all predictors and targets required to replicate the results, are publicly available on Open Science Foundation, at: https://osf.io/rn7qb/?view_only=f2c4b60f7b2e4e80a04151c67e6e6d52

## References

1. Pedersen, P. M., Stig Jørgensen, H., Nakayama, H., Raaschou, H. O., & Olsen, T. S. (1995). Aphasia in acute stroke: incidence, determinants, and recovery. Annals of Neurology: Official Journal of the American Neurological Association and the Child Neurology Society, 38(4), 659–666.

2. Lazar, R. M., Speizer, A. E., Festa, J. R., Krakauer, J. W., & Marshall, R. S. (2008). Variability in language recovery after first-time stroke. Journal of Neurology, Neurosurgery & Psychiatry, 79(5), 530–534.

3. Brady, M. C., Kelly, H., Godwin, J., Enderby, P., & Campbell, P. (2016). Speech and language therapy for aphasia following stroke. Cochrane database of systematic reviews, (6).

4. Iorga M, Higgins J, Caplan D, Zinbarg R, Kiran S, Thompson CK, Rapp B, Parrish TB (2021). Predicting language recovery in post-stroke aphasia using behavior and functional MRI. Sci Rep.;11(1):8419.

5. Marte MJ, Carpenter E, Scimeca M, Russell-Meill M, Peñaloza C, Grasemann U, Miikkulainen R, Kiran S (2025). Machine Learning Predictions of Recovery in Bilingual Poststroke Aphasia: Aligning Insights With Clinical Evidence. Stroke. 56(2):494–504.

6. Stefaniak, J. D., Geranmayeh, F., & Lambon Ralph, M. A. (2022). The multidimensional nature of aphasia recovery post-stroke. Brain, 145(4), 1354–1367.

7. Benghanem S, Rosso C, Arbizu C, Moulton E, Dormont D, Leger A, Pires C, Samson Y (2019). Aphasia outcome: the interactions between initial severity, lesion size and location. J Neurol. 266(6):1303–1309.

8. Wilson SM, Entrup JL, Schneck SM, Onuscheck CF, Levy DF, Rahman M, Willey E, Casilio M, Yen M, Brito AC, Kam W, Davis LT, de Riesthal M, Kirshner HS (2023). Recovery from aphasia in the first year after stroke. Brain. 2023 Mar 1;146(3):1021–1039.

9. Zhao Y, Cox CR, Lambon Ralph MA, Halai AD (2023). Using in vivo functional and structural connectivity to predict chronic stroke aphasia deficits. Brain. 146(5):1950–1962.

10. Saur, D., Ronneberger, O., Kümmerer, D., Mader, I., Weiller, C., & Klöppel, S. (2010). Early functional magnetic resonance imaging activations predict language outcome after stroke. Brain, 133(4), 1252–1264.

11. Billot A, Lai S, Varkanitsa M, Braun EJ, Rapp B, Parrish TB, Higgins J, Kurani AS, Caplan D, Thompson CK, Ishwar P, Betke M, Kiran S (2022). Multimodal Neural and Behavioral Data Predict Response to Rehabilitation in Chronic Poststroke Aphasia. Stroke. 53(5):1606–1614.

12. Hu, X., Varkanitsa, M., Kropp, E., Betke, M., Ishwar, P., & Kiran, S. (2025). Aphasia severity prediction using a multi-modal machine learning approach. NeuroImage, 121300.

13. Saur, D., Lange, R., Baumgaertner, A., Schraknepper, V., Willmes, K., Rijntjes, M., & Weiller, C. (2006). Dynamics of language reorganization after stroke. Brain, 129(6), 1371–1384.

14. Anika Stockert, Max Wawrzyniak, Julian Klingbeil, Katrin Wrede, Dorothee Kemmerer, Gesa Hartwigsen, Christoph P Kaller, Cornelius Weiller, Dorothee Saur, (2020). Dynamics of language reorganization after left temporo-parietal and frontal stroke, Brain, Volume 143, Issue 3, Pages 844–861

15. Jiang Z, Kuhnke P, Stockert A, Wawrzyniak M, Halai A, Saur D, Hartwigsen G (2025). Dynamic reorganization of task-related network interactions in post-stroke aphasia recovery. Brain. 148(10):3563–3575.

16. Siegel JS, Seitzman BA, Ramsey LE, Ortega M, Gordon EM, Dosenbach NUF, Petersen SE, Shulman GL, Corbetta M (2018). Re-emergence of modular brain networks in stroke recovery. Cortex.101:44–59.

17. Halai AD, Woollams AM, Lambon Ralph MA (2018). Predicting the pattern and severity of chronic post-stroke language deficits from functionally-partitioned structural lesions. Neuroimage Clin. 19:1–13.

18. De Baene, W., Duyck, W., Brass, M., & Carreiras, M. (2015). Brain circuit for cognitive control is shared by task and language switching. Journal of cognitive neuroscience, 27(9), 1752–1765.

19. Benjamini, Y., & Hochberg, Y. (1995). Controlling the false discovery rate: a practical and powerful approach to multiple testing. Journal of the Royal statistical society: series B (Methodological), 57(1), 289–300.

20. Levy DF, Entrup JL, Schneck SM, Onuscheck CF, Rahman M, Kasdan A, Casilio M, Willey E, Davis LT, de Riesthal M, Kirshner HS, Wilson SM (2024). Multivariate lesion symptom mapping for predicting trajectories of recovery from aphasia. Brain Commun. 6(1):fcae024.

21. Osa García A, Brambati SM, Brisebois A, Désilets-Barnabé M, Houzé B, Bedetti C, Rochon E, Leonard C, Desautels A, Marcotte K (2020). Predicting Early Post-stroke Aphasia Outcome From Initial Aphasia Severity. Front Neurol. 11:120.

22. Lazar RM, Minzer B, Antoniello D, Festa JR, Krakauer JW, Marshall RS (2010). Improvement in aphasia scores after stroke is well predicted by initial severity. Stroke. 41(7):1485–8.

23. Tilwani D, O’Reilly C, Riccardi N, Shalin VL, den Ouden DB, Fridriksson J, Shinkareva SV, Sheth AP, Desai RH (2025). Benchmarking machine learning models in lesion-symptom mapping for predicting language outcomes in stroke survivors. Front Neuroimaging. 4:1573816.

24. Warren DE, Power JD, Bruss J, Denburg NL, Waldron EJ, Sun H, Petersen SE, Tranel D (2014). Network measures predict neuropsychological outcome after brain injury. Proc Natl Acad Sci U S A. 111(39):14247–52.

25. Tao, Y., & Rapp, B. (2021). Investigating the network consequences of focal brain lesions through comparisons of real and simulated lesions. Scientific Reports, 11(1), 2213.

26. Wawrzyniak, M., Schneider, H. R., Klingbeil, J., Stockert, A., Hartwigsen, G., Weiller, C., & Saur, D. (2022). Resolution of diaschisis contributes to early recovery from post-stroke aphasia. Neuroimage, 251, 119001.

27. Stockert, A., Kümmerer, D., & Saur, D. (2016). Insights into early language recovery: From basic principles to practical applications. Aphasiology, 30(5), 517–541.

28. Billot, A., de Schotten, M. T., Parrish, T. B., Thompson, C. K., Rapp, B., Caplan, D., & Kiran, S. (2022). Structural disconnections associated with language impairments in chronic post-stroke aphasia using disconnectome maps. Cortex, 155, 90–106.

29. Kristinsson, S., Busby, N., Rorden, C., Newman-Norlund, R., Den Ouden, D. B., Magnusdottir, S., … & Fridriksson, J. (2022). Brain age predicts long-term recovery in post-stroke aphasia. Brain Communications, 4(5), fcac252.

30. Stefaniak, J. D., Halai, A. D., & Lambon Ralph, M. A. (2020). The neural and neurocomputational bases of recovery from post-stroke aphasia. Nature Reviews Neurology, 16(1), 43–55.

31. Hartwigsen, G. (2018). Flexible redistribution in cognitive networks. Trends in cognitive sciences, 22(8), 687–698.

32. D’Esposito, M., Deouell, L. Y., & Gazzaley, A. (2003). Alterations in the BOLD fMRI signal with ageing and disease: a challenge for neuroimaging. Nature Reviews Neuroscience, 4(11), 863–872.

33. Duncan, J. (2010). The multiple-demand (MD) system of the primate brain: mental programs for intelligent behaviour. Trends in cognitive sciences, 14(4), 172–179.

34. Geranmayeh, F., Chau, T. W., Wise, R. J., Leech, R., & Hampshire, A. (2017). Domain-general subregions of the medial prefrontal cortex contribute to recovery of language after stroke. Brain, 140(7), 1947–1958.

35. Turkeltaub PE, Martin KC, Laks AB, DeMarco AT. Right hemisphere language network plasticity in aphasia. Brain. 2025 Sep 8:awaf308.

36. Abel S, Weiller C, Huber W, Willmes K, Specht K (2015). Therapy-induced brain reorganization patterns in aphasia. Brain. 138(Pt 4):1097–112.

37. Nenert R, Allendorfer JB, Martin AM, Banks C, Vannest J, Holland SK, Hart KW, Lindsell CJ, Szaflarski JP (2018). Longitudinal fMRI study of language recovery after a left hemispheric ischemic stroke. Restor Neurol Neurosci. 36(3):359–385.

38. Hartwigsen G, Saur D (2019). Neuroimaging of stroke recovery from aphasia - Insights into plasticity of the human language network. Neuroimage.190:14–31.

39. Hartwigsen G, Bzdok D, Klein M, Wawrzyniak M, Stockert A, Wrede K, Classen J, Saur D (2017). Rapid short-term reorganization in the language network. Elife. 6:e25964.

40. Ding, X., Zhang, S., Huang, W., Zhang, S., Zhang, L., Hu, J., Li, J., Ge, Q., Wang, Y., Ye, X., & Zhang, J. (2022). Comparative efficacy of non-invasive brainstimulation for post-stroke aphasia: A networkmeta-analysis and meta-regression of moderators.Neuroscience and Biobehavioral Reviews, 140,104804.

41. Twardochleb, Z., Szczepańska, M., Miś, M. M., Miś, M., Druszcz, A., Paprocka-Borowicz, M., & Rosińczuk, J. (2025). tDCS and speech therapy in aphasia treatment: a multicenter comparative study of efficacy. Medical science monitor: international medical journal of experimental and clinical research, 31, e950237.

42. Ren, J., Ren, W., Zhou, Y., Dahmani, L., Duan, X., Fu, X., … & Liu, H. (2023). Personalized functional imaging-guided rTMS on the superior frontal gyrus for post-stroke aphasia: a randomized sham-controlled trial. Brain Stimulation, 16(5), 1313–1321.

43. Turkeltaub, P. E., Martin, K. C., Laks, A. B., & DeMarco, A. T. (2026). Right hemisphere language network plasticity in aphasia. Brain, 149(4), 1396–1409.

44. Zumbansen A, Black SE, Chen JL, J Edwards D, Hartmann A, Heiss WD, Lanthier S, Lesperance P, Mochizuki G, Paquette C, Rochon EA, Rubi-Fessen I, Valles J, Kneifel H, Wortman-Jutt S, Thiel A; NORTHSTAR-study group (2020). Non-invasive brain stimulation as add-on therapy for subacute post-stroke aphasia: a randomized trial (NORTHSTAR). Eur Stroke J. 5(4):402–413.

45. Hartwigsen G, Neef NE, Camilleri JA, Margulies DS, Eickhoff SB (2019). Functional Segregation of the Right Inferior Frontal Gyrus: Evidence From Coactivation-Based Parcellation. Cereb Cortex. 29(4):1532–1546.

46. Cao Y, Vikingstad EM, George KP, Johnson AF, Welch KM (1999). Cortical language activation in stroke patients recovering from aphasia with functional MRI. Stroke. 30(11):2331–40.

47. Fan L, Li C, Huang ZG, Zhao J, Wu X, Liu T, Li Y, Wang J (2022). The longitudinal neural dynamics changes of whole brain connectome during natural recovery from poststroke aphasia. Neuroimage Clin. 36:103190.

48. Huber W, Poeck K, Willmes K (1984). The Aachen Aphasia Test. Adv Neurol. 42:291–303.

49. Hickok, G., & Poeppel, D. (2007). The cortical organization of speech processing. Nature reviews neuroscience, 8(5), 393–402.

50. Friederici, A. D., & Gierhan, S. M. (2013). The language network. Current opinion in neurobiology, 23(2), 250–254.

51. Fedorenko, E., & Thompson-Schill, S. L. (2014). Reworking the language network. Trends in cognitive sciences, 18(3), 120–126.

52. Basilakos A, Smith KG, Fillmore P, Fridriksson J, Fedorenko E. Functional Characterization of the Human Speech Articulation Network. Cereb Cortex. 2018 May 1;28(5):1816–1830.

53. Combrisson, E., & Jerbi, K. (2015). Exceeding chance level by chance: The caveat of theoretical chance levels in brain signal classification and statistical assessment of decoding accuracy. Journal of neuroscience methods, 250, 126–136.

54. Pereira, F., Mitchell, T., & Botvinick, M. (2009). Machine learning classifiers and fMRI: a tutorial overview. Neuroimage, 45(1), S199–S209.

55. Mumford, J. A., Poline, J. B., & Poldrack, R. A. (2015). Orthogonalization of regressors in fMRI models. PloS one, 10(4), e0126255.

56. Varoquaux, G., Raamana, P. R., Engemann, D. A., Hoyos-Idrobo, A., Schwartz, Y., & Thirion, B. (2017). Assessing and tuning brain decoders: cross-validation, caveats, and guidelines. NeuroImage, 145, 166–179.

57. Covert, I., Lundberg, S., & Lee, S. I. (2021). Explaining by removing: A unified framework for model explanation. Journal of Machine Learning Research, 22(209), 1–90.

