## Supplementary Information for "Changes in Neural Dynamics of Brain Activity and Connectivity Independently Predict Post-Stroke Aphasia Recovery"

**Short title:** Neuroimaging predictors of aphasia recovery

**Word count:** Abstract: 200 words, **Main Text:** < 5000 words

**Number of Figures:** 6

**Contact details**

| Andrea Bruera | Anika Stockert | Gesa Hartwigsen |
|---|---|---|
| | | |

### Supplementary Information (SI)

#### SI Section 1. Details of the fMRI acquisition procedure and experimental paradigm

Stimuli were adopted from a previous study in the visual domain [1SI] and consisted of simple sentences with a highly predictable ending that followed a regular pattern (i.e., a person performing a typical job). As we aimed to analyze as many longitudinal data sets as possible, we combined data from two studies [13, 2SI] and accepted slight differences between paradigms. In both studies, correct (e.g., 'The pilot flies the plane') and temporally reversed versions of the same stimulus elements (e.g., 'enalp eht seilf tolip ehT') were presented auditorily. Firstly, the paradigms differed in terms of another type of stimulus that was excluded from the analysis: Semantically incorrect sentences [9] and pseudo speech [2SI] . The second difference was related to button press instructions that either targeted incorrect or reverse played sentences [13] or each stimulus [2SI] . Lastly, the number of stimuli differed and included 46 correct and 46 reversed sentences distributed across six sessions [13] versus 30 correct and 30 reversed sentences presented in a single session [2SI] . To ensure that task instructions had been understood, prior to in-scanner testing all participants were familiarized with the stimuli and tasks in a training session in which performance had to be above chance to further participate in the experiment.

#### SI Section 2. Functional brain activity

As measures of functional brain activity, we defined a first-level General Linear Model (GLM). Correct and reversed speech were modelled as separate conditions; acute, subacute and chronic phases after the stroke as separate time points. Sentence onsets and durations were modelled as stimulus (boxcar) functions. Head motions were added as regressors of no interest. Signal drifts were accounted for by Discrete Cosine Transform basis (128s cut-off). First (time) and second (dispersion) derivatives were added to the model in order to account for variability in the time-to-peak and potential differences in the shape of the HRF. These event regressors were convolved with the canonical haemodynamic response function (HRF). Finally, we computed the contrast between correct and reversed sentence speech. To obtain the overall measures of brain activity for each subject in each ROI, we averaged the GLM betas for the voxels contained within each subject-specific ROI.

#### SI Section 3. Effective connectivity

To obtain effective connectivity measures during speech processing, we used Dynamic Causal Modelling (DCM) in SPM12 implemented in MATLAB (version 23.2; [3SI] ). A full model was specified and estimated for each participant, which assumed that all ROIs were fully connected via reciprocal connections. We set the onset of speech and reversed speech as modulatory inputs on the connections between the 6 ROIs.

The first eigenvariate of the time series was extracted from the subject-specific ROIs and adjusted for effects-of-interest. We extracted modulatory parameters of speech on the effective connectivity of 9 connections of interest from each individual participant's estimated DCM model. The 9 connections of interest were from multiple-demand regions to undamaged language areas (SMA/dACC to left IFG, SMA/dACC to left PTL, left DLPFC to left IFG, left DLPFC to left PTL, right DLPFC to left IFG, right DLPFC to left PTL), interactions within the language network (left IFG to left PTL, left PTL to left IFG), and from right IFG to left IFG (cf. [15]).

#### SI Section 4. Details on Confound Control

We used the cross-validated confound-control procedure of [28] . First, we used the training data for a linear regression model to learn to predict each target from the lesion affection of the four left hemisphere ROIs. Second, we predicted the targets for the subjects in both the training and the test sets from their lesion affection and age. Finally, we computed the residual values for language ability and improvement (i.e., the difference between the true and the predicted targets), to use them as actual targets for training and testing the actual predictive model.

In addition to lesion affection and age, we also added functional brain activity when using effective connectivity as predictors and *vice versa* for brain activity (e.g., when using acute connectivity measures as predictors, we removed the variance explained by lesion affection, age, *and* acute brain activity). This ensured prediction performance reflected independent contributions of each set of predictors above and beyond all other predictors. Please note that, as described above, for language improvement we additionally added the language comprehension score at a previous time point to the confound variables thereby controlling for aphasia severity as another predictor of outcome.

**SI Figure 1. Brain lesion overlay - Surface-projected lateral and medial view of the left hemisphere and sagittal view of both hemispheres.** Left panel: projection of the left-hemisphere lesion maps for the 41 patients considered in this study on the Freesurfer template surface. Right panel: lesion maps for the 47 patients considered in this study, plotted against the MNI template. The intensity of the shade (in a perceptually uniform palette, roughly from blue to yellow) indicates the amount of participants for whom the underlying voxel was damaged.

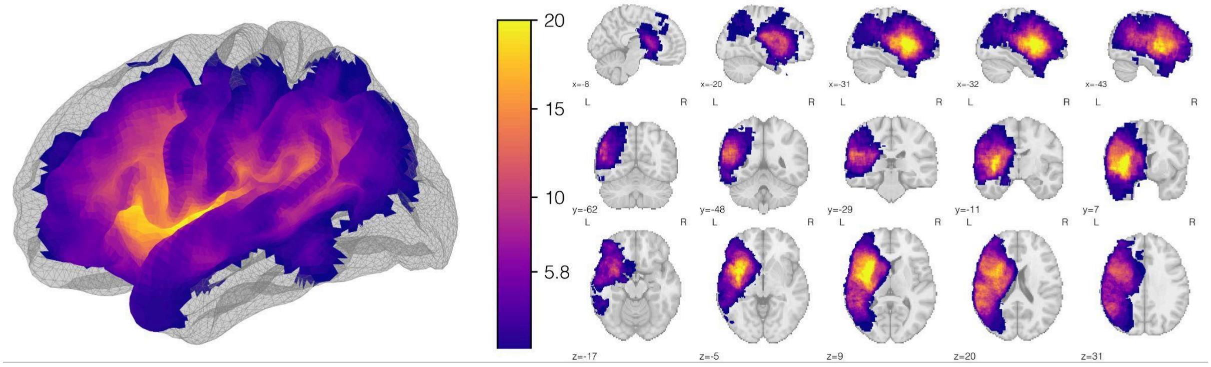


**SI Table 1a. Patient demographic details and language comprehension scores.** AAT= Aachen Aphasia Test. Scores range from 0 to 1, with a score of 1 indicating full recovery.

| *Subject* | *Age* | *Sex* | *Acute AAT comprehension score* | *Subacute AAT comprehension score* | *Chronic AAT comprehension score* | *Average AAT comprehension score* |
|---|---|---|---|---|---|---|
| sub-01 | 69 | M | 0.512 | 0.724 | 0.965 | 0.7337 |
| sub-02 | 63 | M | 0.941 | 0.971 | 0.982 | 0.9647 |
| sub-03 | 69 | M | 0.835 | 0.941 | 0.971 | 0.9157 |
| sub-04 | 55 | M | 0.829 | 0.918 | 0.937 | 0.8947 |
| sub-05 | 72 | F | 0.288 | 0.406 | 0.753 | 0.4823 |
| sub-06 | 40 | F | 0.271 | 0.435 | 0.871 | 0.5257 |
| sub-07 | 64 | M | 0.977 | 1.0 | 1.0 | 0.9923 |
| sub-08 | 58 | M | 0.941 | 1.0 | 1.0 | 0.9803 |
| sub-09 | 59 | M | 0.729 | 0.888 | 0.918 | 0.845 |
| sub-10 | 31 | M | 0.829 | 0.924 | 0.982 | 0.9117 |
| sub-11 | 68 | F | 0.341 | 0.394 | 0.553 | 0.4293 |
| sub-12 | 51 | M | 0.429 | 0.682 | 0.877 | 0.6627 |
| sub-13 | 38 | M | 0.924 | 0.982 | 0.982 | 0.9627 |
| sub-14 | 69 | M | 0.341 | 0.659 | 0.941 | 0.647 |
| sub-15 | 56 | M | 0.012 | 0.082 | 0.494 | 0.196 |
| sub-16 | 71 | M | 0.171 | 0.553 | 0.847 | 0.5237 |
| sub-17 | 61 | F | 0.024 | 0.412 | 0.688 | 0.3747 |
| sub-18 | 34 | M | 0.977 | 0.977 | 0.988 | 0.9807 |

| | | | | | | |
|---|---|---|---|---|---|---|
| sub-19 | 66 | F | 0.029 | 0.671 | 0.953 | 0.551 |
| sub-20 | 71 | M | 0.035 | 0.341 | 0.629 | 0.335 |
| sub-21 | 61 | M | 0.024 | 0.424 | 0.929 | 0.459 |
| sub-22 | 76 | M | 0.371 | 0.559 | 0.747 | 0.559 |
| sub-23 | 55 | M | 0.394 | 0.465 | 0.659 | 0.506 |
| sub-24 | 64 | M | 0.912 | 0.971 | 0.977 | 0.9533 |
| sub-25 | 73 | F | 0.035 | 0.035 | 0.488 | 0.186 |
| sub-26 | 61 | M | 0.129 | 0.588 | 0.818 | 0.5117 |
| sub-27 | 44 | F | 0.8 | 0.971 | 0.982 | 0.9177 |
| sub-28 | 68 | M | 0.6 | 0.788 | 0.853 | 0.747 |
| sub-29 | 65 | M | 0.4 | 0.447 | 0.894 | 0.5803 |
| sub-30 | 58 | F | 0.447 | 0.765 | 0.918 | 0.71 |
| sub-31 | 55 | M | 0.147 | 0.4 | 0.759 | 0.4353 |
| sub-32 | 65 | M | 0.465 | 0.912 | 0.959 | 0.7787 |
| sub-33 | 55 | M | 0.518 | 0.647 | 0.9 | 0.6883 |
| sub-34 | 37 | M | 0.759 | 0.918 | 0.965 | 0.8807 |
| sub-35 | 57 | M | 0.865 | 0.929 | 0.959 | 0.9177 |
| sub-36 | 16 | M | 0.424 | 0.871 | 1.0 | 0.765 |
| sub-37 | 43 | M | 0.0 | 0.482 | 0.835 | 0.439 |
| sub-38 | 39 | F | 0.0 | 0.365 | 0.518 | 0.2943 |
| sub-39 | 64 | F | 0.3 | 0.335 | 0.565 | 0.4 |
| sub-40 | 68 | M | 0.465 | 0.6 | 0.653 | 0.5727 |
| sub-41 | 26 | F | 0.547 | 0.547 | 0.994 | 0.696 |
| sub-42 | 29 | M | 0.294 | 0.294 | 0.882 | 0.49 |
| sub-43 | 52 | M | 0.853 | 0.853 | 0.947 | 0.8843 |
| sub-44 | 40 | F | 0.806 | 0.806 | 0.906 | 0.8393 |
| sub-45 | 50 | F | 0.382 | 0.382 | 0.718 | 0.494 |
| sub-46 | 49 | M | 0.582 | 0.582 | 0.829 | 0.6643 |
| sub-47 | 61 | M | 0.6 | 0.6 | 0.706 | 0.6353 |

**SI Table 1b. Patient details of lesion affection (percentage of damaged voxels) in each of the four ROIs used for the analyses.**

| *Subject* | *Lesion % left DLPFC* | *Lesion % left IFG* | *Lesion % left PTL* | *lesion % left SMA* |
|---|---|---|---|---|
| sub-01 | 59.76% | 1.14% | 10.07% | 0.0% |
| sub-02 | 2.06% | 0.0% | 30.39% | 0.0% |
| sub-03 | 0.0% | 0.0% | 0.57% | 0.0% |
| sub-04 | 0.0% | 0.0% | 1.42% | 0.0% |

| | | | | |
|---|---|---|---|---|
| sub-05 | 71.91% | 23.84% | 0.0% | 0.52% |
| sub-06 | 7.27% | 9.6% | 0.28% | 0.0% |
| sub-07 | 30.59% | 0.21% | 0.0% | 0.0% |
| sub-08 | 0.0% | 0.0% | 13.01% | 0.0% |
| sub-09 | 0.0% | 0.0% | 23.46% | 0.0% |
| sub-10 | 0.0% | 0.0% | 34.38% | 0.0% |
| sub-11 | 32.0% | 13.31% | 0.0% | 0.0% |
| sub-12 | 10.95% | 29.62% | 0.0% | 0.0% |
| sub-13 | 0.0% | 0.0% | 8.55% | 0.0% |
| sub-14 | 68.0% | 59.13% | 0.38% | 0.0% |
| sub-15 | 99.57% | 91.43% | 42.45% | 3.04% |
| sub-16 | 0.65% | 61.3% | 24.22% | 0.0% |
| sub-17 | 31.13% | 14.76% | 15.0% | 0.0% |
| sub-18 | 0.65% | 0.0% | 5.41% | 0.0% |
| sub-19 | 87.85% | 49.64% | 9.02% | 23.13% |
| sub-20 | 8.46% | 26.01% | 26.59% | 0.0% |
| sub-21 | 60.85% | 23.01% | 0.0% | 0.0% |
| sub-22 | 0.0% | 0.0% | 37.51% | 0.0% |
| sub-23 | 0.76% | 9.8% | 98.67% | 0.0% |
| sub-24 | 0.0% | 0.0% | 21.75% | 0.0% |
| sub-25 | 70.93% | 72.65% | 0.76% | 0.43% |
| sub-26 | 0.0% | 1.03% | 28.77% | 0.0% |
| sub-27 | 0.87% | 23.01% | 0.0% | 0.0% |
| sub-28 | 0.0% | 0.0% | 10.16% | 0.0% |
| sub-29 | 49.02% | 0.93% | 0.0% | 0.0% |
| sub-30 | 0.11% | 1.14% | 0.0% | 0.0% |
| sub-31 | 6.62% | 78.74% | 23.55% | 0.0% |
| sub-32 | 0.54% | 0.0% | 2.75% | 0.0% |
| sub-33 | 0.0% | 0.0% | 62.87% | 0.0% |
| sub-34 | 0.0% | 0.0% | 35.71% | 0.0% |
| sub-35 | 0.0% | 0.0% | 11.3% | 0.0% |
| sub-36 | 43.06% | 11.56% | 0.0% | 0.0% |
| sub-37 | 1.84% | 0.0% | 37.8% | 0.0% |
| sub-38 | 2.49% | 15.79% | 30.48% | 4.78% |
| sub-39 | 0.11% | 2.89% | 1.99% | 0.0% |
| sub-40 | 4.34% | 39.01% | 7.88% | 0.0% |
| sub-41 | 92.3% | 12.18% | 6.36% | 0.0% |

| | | | | |
|---|---|---|---|---|
| sub-42 | 0.22% | 11.56% | 27.64% | 0.0% |
| sub-43 | 0.22% | 30.24% | 0.0% | 0.0% |
| sub-44 | 0.0% | 0.0% | 10.92% | 0.0% |
| sub-45 | 56.83% | 65.94% | 0.0% | 0.0% |
| sub-46 | 0.0% | 0.0% | 60.11% | 0.0% |
| sub-47 | 0.0% | 0.0% | 27.73% | 0.0% |

**SI Figure 2. Ridge regression prediction scores when using lesion affection, age and previous language ability as predictors.** Scores refer to the prediction performance after removing the variance explained by the other confounds (i.e. age/lesions/previous ability) as well as the neuroimaging predictors analyzed in the main paper (activity and connectivity) at the corresponding time point.

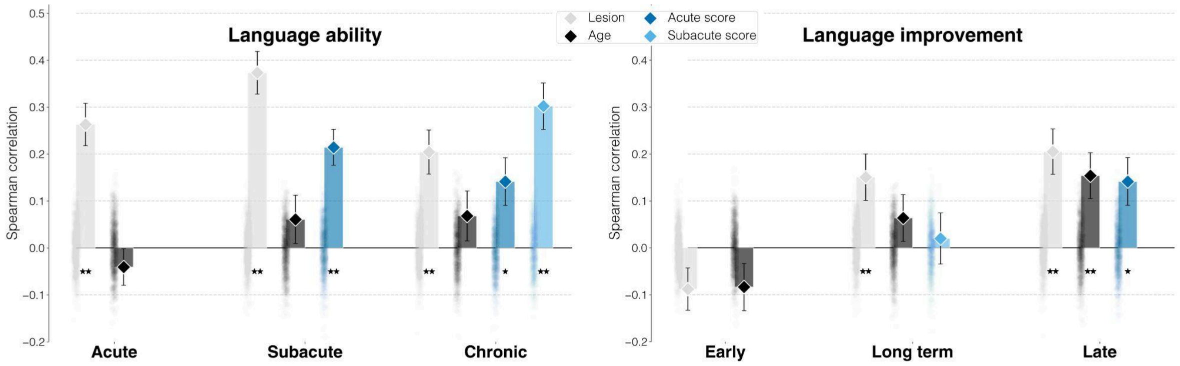


**SI Table 2a-b. Detailed statistics for SI Figure 2.** Bold font indicates statistical significance after FDR correction.

**SI Table 2a**

| *Predictor* | *Target* | *Spearman correlation* | *FDR-corrected p-value* | *Raw p-value* |
|---|---|---|---|---|
| **Lesions** | **Acute ability** | **0.263** | **0.0024** | **0.001** |
| **Lesions** | **Subacute ability** | **0.373** | **0.0024** | **0.001** |
| **Lesions** | **Chronic ability** | **0.204** | **0.0024** | **0.001** |
| Age | Acute ability | -0.041 | 0.8684 | 0.7662 |
| Age | Subacute ability | 0.061 | 0.1458 | 0.1029 |
| Age | Chronic ability | 0.068 | 0.1436 | 0.0929 |
| **Acute score** | **Subacute ability** | **0.214** | **0.0024** | **0.001** |
| **Acute score** | **Chronic ability** | **0.141** | **0.0085** | **0.005** |
| **Subacute score** | **Chronic ability** | **0.302** | **0.0024** | **0.001** |

**SI Table 2b**

| *Predictor* | *Target* | *Spearman correlation* | *FDR-corrected p-value* | *Raw p-value* |
|---|---|---|---|---|

| | | | | |
|---|---|---|---|---|
| Lesions | Acute to subacute improvement | -0.088 | 0.964 | 0.964 |
| **Lesions** | **Acute to chronic improvement** | **0.151** | **0.0024** | **0.001** |
| **Lesions** | **Subacute to chronic improvement** | **0.205** | **0.0024** | **0.001** |
| Age | Acute to subacute improvement | -0.084 | 0.964 | 0.959 |
| Age | Acute to chronic improvement | 0.064 | 0.1646 | 0.1259 |
| **Age** | **Subacute to chronic improvement** | **0.154** | **0.0042** | **0.002** |
| **Acute score** | **Subacute to chronic improvement** | **0.141** | **0.0057** | **0.003** |
| Subacute score | Acute to chronic improvement | 0.02 | 0.4513 | 0.3716 |

**SI Table 3a-b. Full set of results and statistics for the main analyses - Figure 3 (main text).** Bold font indicates statistical significance after FDR correction. Acute to subacute improvement=early improvement, acute to chronic improvement=long-term improvement, subacute to chronic improvement=late improvement.

**SI Table 3a**

| *Predictor* | *Target* | *Spearman correlation* | *FDR-corrected p-value* | *Raw p-value* |
|---|---|---|---|---|
| **Acute activity** | **Acute ability** | **0.163** | **0.0028** | **0.001** |
| **Acute activity** | **Subacute ability** | **0.143** | **0.0043** | **0.002** |
| **Subacute activity** | **Subacute ability** | **0.366** | **0.0028** | **0.001** |
| **Acute activity** | **Chronic ability** | **0.235** | **0.0028** | **0.001** |
| **Subacute activity** | **Chronic ability** | **0.286** | **0.0028** | **0.001** |
| Chronic activity | Chronic ability | -0.085 | 0.968 | 0.957 |
| Acute connectivity | Acute ability | -0.045 | 0.9522 | 0.8162 |
| Acute connectivity | Subacute ability | 0.055 | 0.2253 | 0.1449 |
| **Subacute connectivity** | **Subacute ability** | **0.189** | **0.0043** | **0.002** |

| | | | | |
|---|---|---|---|---|
| Acute connectivity | Chronic ability | 0.066 | 0.1382 | 0.0839 |
| Subacute connectivity | Chronic ability | 0.022 | 0.4234 | 0.3327 |
| Chronic connectivity | Chronic ability | -0.009 | 0.7042 | 0.5784 |

**SI Table 3b**

| | | | | |
|---|---|---|---|---|
| **Acute activity** | **Acute to subacute improvement** | **0.208** | **0.0028** | **0.001** |
| **Subacute activity** | **Acute to subacute improvement** | **0.312** | **0.0028** | **0.001** |
| **Acute activity** | **Acute to chronic improvement** | **0.123** | **0.0168** | **0.009** |
| **Subacute activity** | **Acute to chronic improvement** | **0.219** | **0.0028** | **0.001** |
| **Chronic activity** | **Acute to chronic improvement** | **0.18** | **0.0028** | **0.001** |
| Acute activity | Subacute to chronic improvement | 0.04 | 0.3105 | 0.2218 |
| **Subacute activity** | **Subacute to chronic improvement** | **0.141** | **0.0043** | **0.002** |
| **Chronic activity** | **Subacute to chronic improvement** | **0.116** | **0.016** | **0.008** |
| **Acute connectivity** | **Acute to subacute improvement** | **0.257** | **0.0028** | **0.001** |
| **Subacute connectivity** | **Acute to subacute improvement** | **0.211** | **0.0028** | **0.001** |
| Acute connectivity | Acute to chronic improvement | -0.072 | 0.968 | 0.9191 |
| Subacute connectivity | Acute to chronic improvement | 0.083 | 0.0717 | 0.041 |
| Chronic connectivity | Acute to chronic improvement | 0.045 | 0.2267 | 0.1538 |
| Acute connectivity | Subacute to chronic improvement | -0.091 | 0.968 | 0.968 |
| Subacute connectivity | Subacute to chronic improvement | -0.076 | 0.968 | 0.9331 |
| Chronic connectivity | Subacute to chronic improvement | 0.03 | 0.3823 | 0.2867 |

**SI Table 4a-c. Detailed statistics for Figure 4 (main text) - brain activity predicts chronic language ability and improvement.** Bold font indicates statistical significance for the directionality of the effect, if the impact was negative (i.e.

removing the predictor decreased prediction performance), after FDR correction. Acute to subacute improvement=early improvement, acute to chronic improvement=long-term improvement, subacute to chronic improvement=late improvement.

**SI Table 4a**

| *Predictors* | *Removed predictor* | *Target* | *Spearman correlation* | *FDR-corrected p-value* | *Average weight* | *Percentage decrease* |
|---|---|---|---|---|---|---|
| Acute activity | L DLPFC | Acute ability | 0.1293 | 0.0052 | 0.0247 | -20.5% |
| **Acute activity** | **L DLPFC** | **Subacute ability** | **0.0273** | **0.2927** | **0.0531** | **-80.8%** |
| **Acute activity** | **L DLPFC** | **Chronic ability** | **0.0723** | **0.0957** | **0.0382** | **-69.2%** |
| **Acute activity** | **L DLPFC** | **Early improvement** | **0.044** | **0.2081** | **0.0522** | **-78.8%** |
| **Acute activity** | **L DLPFC** | **Longterm improvement** | **-0.0857** | **0.955** | **0.0144** | **-169.8%** |
| **Acute activity** | **L SMA** | **Acute ability** | **0.0827** | **0.0522** | **0.0308** | **-49.2%** |
| Acute activity | L SMA | Subacute ability | 0.1253 | 0.0105 | 0.0335 | -12.1% |
| Acute activity | L SMA | Chronic ability | 0.2217 | 0.0015 | 0.0125 | -5.7% |
| Acute activity | L SMA | Early improvement | 0.2353 | 0.0015 | 0.0001 | 13.3% |
| Acute activity | L SMA | Longterm improvement | 0.1297 | 0.0094 | 0.0002 | 5.7% |
| Acute activity | L IFGorb | Acute ability | 0.1657 | 0.0015 | 0.0257 | 1.8% |
| Acute activity | L IFGorb | Subacute ability | 0.1977 | 0.0015 | 0.0378 | 38.6% |
| Acute activity | L IFGorb | Chronic ability | 0.2697 | 0.0015 | 0.0026 | 14.8% |
| Acute activity | L IFGorb | Early improvement | 0.2453 | 0.0015 | 0.0094 | 18.1% |
| Acute activity | L IFGorb | Longterm improvement | 0.134 | 0.0085 | -0.0049 | 9.2% |
| Acute activity | L PTL | Acute ability | 0.1657 | 0.0015 | 0.004 | 1.8% |
| Acute activity | L PTL | Subacute ability | 0.1523 | 0.0028 | -0.0309 | 6.8% |
| Acute activity | L PTL | Chronic ability | 0.238 | 0.0015 | -0.0071 | 1.3% |
| Acute activity | L PTL | Early improvement | 0.159 | 0.0015 | -0.0572 | -23.4% |
| Acute activity | L PTL | Longterm improvement | 0.162 | 0.0015 | -0.0035 | 32.1% |
| Acute activity | R DLPFC | Acute ability | 0.1893 | 0.0015 | -0.0009 | 16.4% |
| Acute activity | R DLPFC | Subacute | 0.1267 | 0.0094 | -0.0343 | -11.2% |

| | | | | | | |
|---|---|---|---|---|---|---|
| | | ability | | | | |
| Acute activity | R DLPFC | Chronic ability | 0.2243 | 0.0015 | -0.0092 | -4.5% |
| Acute activity | R DLPFC | Early improvement | 0.1613 | 0.0015 | -0.0587 | -22.3% |
| Acute activity | R DLPFC | Longterm improvement | 0.1553 | 0.0028 | -0.0058 | 26.6% |
| Acute activity | R IFGorb | Acute ability | 0.2017 | 0.0015 | -0.018 | 24.0% |
| Acute activity | R IFGorb | Subacute ability | 0.245 | 0.0015 | -0.019 | 71.7% |
| Acute activity | R IFGorb | Chronic ability | 0.288 | 0.0015 | -0.0041 | 22.6% |
| Acute activity | R IFGorb | Early improvement | 0.1863 | 0.0015 | 0.0169 | -10.3% |
| Acute activity | R IFGorb | Longterm improvement | 0.1507 | 0.0052 | 0.0041 | 22.8% |

**SI Table 4b**

| *Predictors* | *Removed predictor* | *Target* | *Spearman correlation* | *FDR-corrected p-value* | *Average weight* | *Percentage decrease* |
|---|---|---|---|---|---|---|
| Subacute activity | L DLPFC | Subacute ability | 0.333 | 0.0015 | 0.0632 | -9.0% |
| Subacute activity | L DLPFC | Chronic ability | 0.2877 | 0.0015 | 0.0224 | 0.6% |
| Subacute activity | L DLPFC | Early improvement | 0.3173 | 0.0015 | -0.0046 | 1.7% |
| Subacute activity | L DLPFC | Longterm improvement | 0.2277 | 0.0015 | -0.0002 | 4.1% |
| Subacute activity | L DLPFC | Late improvement | 0.1297 | 0.0052 | -0.002 | -7.8% |
| Subacute activity | L SMA | Subacute ability | 0.314 | 0.0015 | 0.088 | -14.2% |
| Subacute activity | L SMA | Chronic ability | 0.1147 | 0.0146 | 0.0358 | -59.9% |
| Subacute activity | L SMA | Early improvement | 0.1387 | 0.0052 | 0.0585 | -55.6% |
| Subacute activity | L SMA | Longterm improvement | 0.126 | 0.0074 | 0.0263 | -42.4% |
| Subacute activity | L SMA | Late improvement | 0.1113 | 0.0125 | 0.0026 | -20.9% |
| Subacute activity | L IFGorb | Subacute ability | 0.3603 | 0.0015 | -0.0571 | -1.5% |
| Subacute activity | L IFGorb | Chronic ability | 0.2933 | 0.0015 | -0.0162 | 2.6% |

| | | | | | | |
|---|---|---|---|---|---|---|
| Subacute activity | L IFGorb | Early improvement | 0.2817 | 0.0015 | -0.052 | -9.7% |
| Subacute activity | L IFGorb | Longterm improvement | 0.248 | 0.0015 | -0.0183 | 13.4% |
| Subacute activity | L IFGorb | Late improvement | 0.1623 | 0.0015 | 0.0033 | 15.4% |
| Subacute activity | L PTL | Subacute ability | 0.3967 | 0.0015 | 0.0122 | 8.4% |
| Subacute activity | L PTL | Chronic ability | 0.285 | 0.0015 | 0.0135 | -0.3% |
| Subacute activity | L PTL | Early improvement | 0.3063 | 0.0015 | -0.0118 | -1.8% |
| Subacute activity | L PTL | Longterm improvement | 0.2133 | 0.0015 | 0.0048 | -2.4% |
| Subacute activity | L PTL | Late improvement | 0.1203 | 0.0063 | 0.0052 | -14.5% |
| Subacute activity | R DLPFC | Subacute ability | 0.364 | 0.0015 | 0.0171 | -0.5% |
| Subacute activity | R DLPFC | Chronic ability | 0.2783 | 0.0015 | 0.0025 | -2.7% |
| Subacute activity | R DLPFC | Early improvement | 0.2983 | 0.0015 | 0.0031 | -4.4% |
| Subacute activity | R DLPFC | Longterm improvement | 0.2253 | 0.0015 | -0.0057 | 3.0% |
| Subacute activity | R DLPFC | Late improvement | 0.1673 | 0.0015 | -0.0024 | 19.0% |
| Subacute activity | R IFGorb | Subacute ability | 0.3363 | 0.0015 | -0.0506 | -8.1% |
| Subacute activity | R IFGorb | Chronic ability | 0.289 | 0.0015 | 0.0022 | 1.0% |
| Subacute activity | R IFGorb | Early improvement | 0.354 | 0.0015 | 0.0105 | 13.5% |
| Subacute activity | R IFGorb | Longterm improvement | 0.1973 | 0.0015 | 0.0235 | -9.8% |
| **Subacute activity** | **R IFGorb** | **Late improvement** | **0.0797** | **0.0604** | **0.0144** | **-43.4%** |

**SI Table 4c**

| *Predictors* | *Removed predictor* | *Target* | *Spearman correlation* | *FDR-corrected p-value* | *Average weight* | *Percentage decrease* |
|---|---|---|---|---|---|---|
| Chronic activity | L DLPFC | Longterm improvement | 0.1783 | 0.0015 | 0.0026 | -1.1% |

| | | | | | | |
|---|---|---|---|---|---|---|
| Chronic activity | L DLPFC | Late improvement | 0.1103 | 0.0125 | 0.0025 | -4.9% |
| Chronic activity | L SMA | Longterm improvement | 0.2123 | 0.0015 | -0.0059 | 17.7% |
| Chronic activity | L SMA | Late improvement | 0.144 | 0.0028 | -0.0015 | 24.1% |
| Chronic activity | L IFGorb | Longterm improvement | 0.134 | 0.0063 | 0.0208 | -25.7% |
| **Chronic activity** | **L IFGorb** | **Late improvement** | **0.0557** | **0.1202** | **0.0051** | **-52.0%** |
| Chronic activity | L PTL | Longterm improvement | 0.1687 | 0.0015 | 0.0051 | -6.5% |
| Chronic activity | L PTL | Late improvement | 0.1147 | 0.0094 | -0.0001 | -1.1% |
| Chronic activity | R DLPFC | Longterm improvement | 0.2007 | 0.0015 | -0.0029 | 11.3% |
| Chronic activity | R DLPFC | Late improvement | 0.156 | 0.0015 | -0.0006 | 34.5% |
| **Chronic activity** | **R IFGorb** | **Longterm improvement** | **0.0707** | **0.088** | **0.018** | **-60.8%** |
| **Chronic activity** | **R IFGorb** | **Late improvement** | **0.0437** | **0.1853** | **0.003** | **-62.4%** |

**SI Table 5. Detailed statistics for Figure 5 (main text) - brain connectivity predicts language ability and improvement.** Bold font indicates statistical significance for the directionality of the effect, if the impact was negative (i.e. removing the predictor decreased prediction performance), after FDR correction. Acute to subacute improvement=early improvement, acute to chronic improvement=long-term improvement, subacute to chronic improvement=late improvement.

| *Predictors* | *Removed predictor* | *Target* | *Spearman correlation* | *FDR-corrected p-value* | *Average weight* | *Percentage decrease* |
|---|---|---|---|---|---|---|
| Acute connectivity | L DLPFC to L IFGorb | Early improvement | 0.2286 | 0.0015 | 0.0152 | -11.1% |
| Acute connectivity | L DLPFC to L PTL | Early improvement | 0.2824 | 0.0015 | -0.0025 | 9.9% |
| Acute connectivity | L SMA to L IFGorb | Early improvement | 0.2113 | 0.0015 | 0.0038 | -17.8% |
| Acute connectivity | L SMA to L PTL | Early improvement | 0.2712 | 0.0015 | -0.0065 | 5.5% |
| Acute connectivity | L IFGorb to L PTL | Early improvement | 0.2687 | 0.0015 | -0.0025 | 4.5% |
| Acute connectivity | L PTL to L IFGorb | Early improvement | 0.2833 | 0.0015 | 0.0127 | 10.2% |

| | | | | | | |
|---|---|---|---|---|---|---|
| Acute connectivity | R DLPFC to L IFGorb | Early improvement | 0.2016 | 0.0015 | 0.0222 | -21.6% |
| Acute connectivity | R DLPFC to L PTL | Early improvement | 0.2178 | 0.0015 | -0.0281 | -15.3% |
| Acute connectivity | R IFGorb to L IFGOrb | Early improvement | 0.2081 | 0.0015 | 0.0046 | -19.0% |

**SI Table 5b**

| *Predictors* | *Removed predictor* | *Target* | *Spearman correlation* | *FDR-corrected p-value* | *Average weight* | *Percentage decrease* |
|---|---|---|---|---|---|---|
| **Subacute connectivity** | **L DLPFC to L IFGorb** | **Subacute ability** | **0.0307** | **0.2773** | **0.1276** | **-83.8%** |
| **Subacute connectivity** | **L DLPFC to L IFGorb** | **Early improveme nt** | **0.0444** | **0.1841** | **0.1144** | **-78.9%** |
| Subacute connectivity | L DLPFC to L PTL | Subacute ability | 0.222 | 0.0015 | -0.053 | 17.2% |
| Subacute connectivity | L DLPFC to L PTL | Early improvement | 0.229 | 0.0015 | -0.0383 | 8.7% |
| Subacute connectivity | L SMA to L IFGorb | Subacute ability | 0.2622 | 0.0015 | 0.0013 | 38.5% |
| Subacute connectivity | L SMA to L IFGorb | Early improvement | 0.1778 | 0.0015 | 0.0044 | -15.6% |
| Subacute connectivity | L SMA to L PTL | Subacute ability | 0.2365 | 0.0015 | 0.0327 | 24.9% |
| Subacute connectivity | L SMA to L PTL | Early improvement | 0.1916 | 0.0015 | 0.0761 | -9.0% |
| Subacute connectivity | L IFGorb to L PTL | Subacute ability | 0.1994 | 0.0015 | 0.0142 | 5.3% |
| Subacute connectivity | L IFGorb to L PTL | Early improvement | 0.2529 | 0.0015 | 0.0149 | 20.1% |
| Subacute connectivity | L PTL to L IFGorb | Subacute ability | 0.1286 | 0.0063 | -0.1194 | -32.1% |
| Subacute connectivity | L PTL to L IFGorb | Early improvement | 0.1867 | 0.0015 | -0.052 | -11.3% |
| Subacute connectivity | R DLPFC to L IFGorb | Subacute ability | 0.1627 | 0.0041 | 0.067 | -14.1% |
| Subacute connectivity | R DLPFC to L IFGorb | Early improvement | 0.2662 | 0.0015 | 0.0095 | 26.4% |
| Subacute connectivity | R DLPFC to L PTL | Subacute ability | 0.1937 | 0.0028 | -0.1134 | 2.3% |
| Subacute connectivity | R DLPFC to L PTL | Early improvement | 0.2844 | 0.0015 | -0.0043 | 35.1% |

| | | | | | | |
|---|---|---|---|---|---|---|
| Subacute connectivity | R IFGorb to L IFGOrb | Subacute ability | 0.327 | 0.0015 | 0.0382 | 72.7% |
| Subacute connectivity | R IFGorb to L IFGOrb | Early improvement | 0.1752 | 0.0015 | -0.0491 | -16.8% |

#### Supplementary Information References


1SI. Baumgaertner A, Weiller C, Buchel C (2002). Event-related fMRI reveals cortical sites involved in contextual sentence integration. Neuroimage. 16: 736–45.

2SI. Saur D, Kreher BW, Schnell S, Kümmerer D, Kellmeyer P, Vry MS, Umarova R, Musso M, Glauche V, Abel S, Huber W, Rijntjes M, Hennig J, Weiller C (2008). Ventral and dorsal pathways for language. Proc Natl Acad Sci U S A. 105(46):18035-40.

3SI. Zeidman P, Jafarian A, Corbin N, Seghier ML, Razi A, Price CJ, Friston KJ (2019). A guide to group effective connectivity analysis, part 1: First level analysis with DCM for fMRI. Neuroimage. 200:174-190.
